# Social attention and coordinated movement produce dissociable spectral and network changes during dyadic improvisational dance

**DOI:** 10.64898/2026.09.24.754139

**Authors:** Rachel M. DeLauder, Noor Tasnim, Mackenzie Aychman, Mary Gahagan, Jessica Purevtugs, Jesse Newpol, Ryan Frank, Grace Grizzell, Grace Nobriga, Sarah Rose, Julia C. Basso

## Abstract

Dance is increasingly delivered as an intervention for social and affective functioning, but a live dyadic encounter confounds several distinct ingredients — social attention, verbal engagement, coordinated movement, and in-the-moment generation — so its neural effects cannot be assigned to any one of them. We recorded dual-brain mobile EEG from adult human participants of both sexes (16 female, 2 male) and a dance instructor across six five-minute conditions that vary these ingredients independently: solitary rest, direct eye gaze, conversation, teacher-led and participant-led mirroring, and free improvisation, with concurrent head-mounted accelerometry. Referencing each condition to eye gaze — socially engaging but nearly motionless — rather than to solitary rest partitioned the response. The components proved dissociable. Nonverbal social attention alone reorganized the power spectrum, flattening the aperiodic exponent (g = −0.97) and desynchronizing alpha-band oscillations (g = −0.90), while leaving inter-regional connectivity unchanged (amplitude g = +0.15; phase g = +0.11). Verbal engagement added high-frequency amplitude coupling (g = +2.61) without further spectral change. Coordinated movement added a decrease in theta/alpha coupling involving the frontoparietal-control network (g = −0.59), the signature of an externally directed state, alongside a further spectral shift (g = −0.61). Movement-vigor covariates, corticokinematic coherence, and aperiodic-adjusted power indicated that the spectral effects were cortical rather than myogenic. Four weeks of improvisational dance training increased empathic responsiveness to others’ distress relative to a movie control (g = 0.82). Coordinated social movement thus acts on the brain componentially, with mutual attention rather than motor load establishing the engaged state.

## INTRODUCTION

Studies of social interaction typically treat it as a unitary condition, contrasted against a solitary baseline. An encounter, however, comprises attention to a partner, speech, movement, and the real-time coordination of that movement. In naturalistic paradigms these occur together, so neural changes attributed to social interaction cannot be assigned to any one of them — and paradigms that isolate a single ingredient typically do so by removing the others, leaving a still, silent participant observing a recording(Schilbach et al., 2013). Which component of a real interaction produces the cortical state that accompanies it is therefore unknown. Dance offers a way to ask: it is sustained, whole-body, and inherently reciprocal, and it decomposes naturally into conditions that vary attention, speech, and movement independently. It is also already delivered as an intervention for conditions ranging from psychiatric disorders to neurodegenerative disease(Wu et al., 2022; Zhang and Wei, 2024), so what the brain does during it matters practically as well.

What is known about dance and the brain comes almost entirely from expertise studies. Long-term dance training is associated with differences in sensorimotor and callosal pathways and in regions supporting social function(Hänggi et al., 2010; Giacosa et al., 2016), with structural asymmetry tracking empathy scores in trained dancers(Li et al., 2026). These findings describe the accumulated correlates of years of practice. They cannot say what happens in a brain during dance, or whether a short course of training produces comparable change — the questions that matter most for anyone considering dance as an intervention. The reason for the gap is straightforward: dance is whole-body, weight-shifting, and usually social, and has therefore been inaccessible to the imaging methods used to study it. Mobile electroencephalography has begun to change this(Theofanopoulou et al., 2024), making it feasible to record what the brain does while two people actually move together.

The Synchronicity Hypothesis of Dance makes a claim about that state: within a single brain, dance coordinates activity across seven neurobehavioral systems — sensory, motor, cognitive, social, emotional, rhythmic, and creative(Basso et al., 2020). At the network level this admits two answers: a generalized increase in communication across systems, or a selective reorganization in which externally directed systems couple more tightly while internally oriented systems disengage — the task-positive/task-negative organization described for solitary tasks(Fox et al., 2005), with the frontoparietal control network arbitrating between them(Yin et al., 2022). Two spectral properties index the accompanying state: the aperiodic (1/f) exponent, which tracks arousal(Lendner et al., 2020) and is thought to reflect excitation–inhibition balance(Gao et al., 2017), and alpha-band activity, whose sensorimotor (mu) form desynchronizes during both execution and observation of another’s movement(Oberman et al., 2007). Neither has been measured during real, reciprocal, whole-body interaction.

We decomposed a dyadic encounter into six five-minute conditions: a resting baseline; eye gaze and conversation, socially demanding but nearly still; and teacher-led mirroring, participant-led mirroring, and free improvisation, which add whole-body movement. Improvisational dance served as both the intervention and the condition expected to tax this state most: it cannot be rehearsed, so each moment must be generated while tracking a partner doing the same, and it requires no prior training(Chappell et al., 2021). Mirroring preserves partner-tracking while removing the generative demand, so mirroring versus improvisation isolates in-the-moment generation from interpersonal coordination. Accelerometry was recorded throughout so movement could be measured rather than assumed.

This pilot study therefore examined how coordinated social movement shapes individual brain function, using mobile hyperscanning EEG recorded before and after the intervention. Aim 1 tested whether interactive movement reorganizes individual-brain state, and whether this reflects a generalized increase in cortical communication or a selective task-positive/task-negative reconfiguration, indexed by the aperiodic exponent, alpha-band power, and coupling within and between attentional and frontoparietal-control networks. Aim 2 tested whether any such change is separable from movement and muscle artifact, using accelerometry as a covariate and corticokinematic coherence as an independent probe. Aim 3 tested whether four weeks of training improves socio-emotional functioning. Inter-brain coupling between partners, the complementary dyadic level of this paradigm, is reported separately.

## METHODS

### Participants and study design

Participants were 18 adults recruited through convenience sampling via community outreach, including social media, websites advertising dance classes, and flyers distributed at local businesses throughout the New River Valley. Inclusion criteria were age ≥ 18 years, ability to engage in physical activity, and English proficiency; participants were excluded if they were pregnant, non-ambulatory, or had an untreated neuropsychiatric or neurological condition. All participants had either no or limited dance experience before the study. Participants were randomly assigned to one of two interventions: improvisational dance training or a dance-themed movie-watching control, each comprising two 90-minute sessions per week over four weeks (eight sessions; 12 hours total; **Figure 1**). This protocol was approved by the Virginia Tech Institutional Review Board (#21-798), and all participants provided written informed consent in accordance with the Declaration of Helsinki. Before and after the intervention, all participants completed a battery of self-report questionnaires (see *Behavioral measures*) and underwent dual-brain hyperscanning EEG. This pilot study was not prospectively registered. Analyses of the within-session condition effects were specified before the neural data were examined; the intervention contrasts are reported as exploratory effect-size estimates.

**Figure 1.**
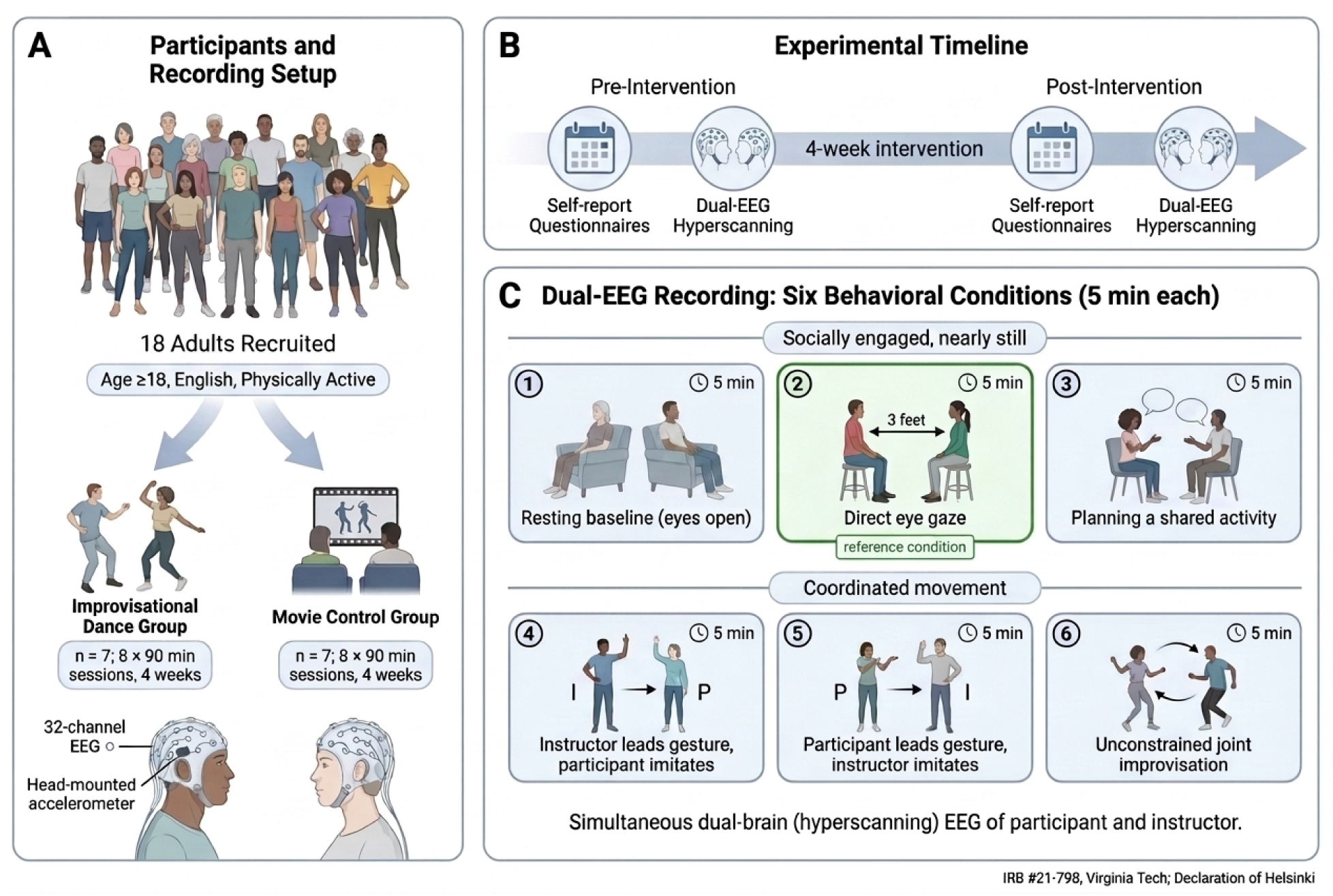
Study design. (**A**) Eighteen adults were recruited and randomized to improvisational dance training (n = 7) or a dance-themed movie-watching control (n = 7), each comprising two 90-minute sessions per week over four weeks. Dual-brain EEG was recorded from participant and instructor with 32-channel mobile systems and head-mounted accelerometry. (**B**) Self-report questionnaires and dual-brain EEG were collected before and after the intervention. (**C**) Six five-minute conditions were recorded in fixed order. Condition 1 was a solitary resting baseline; conditions 2 and 3 are socially engaged but nearly still; conditions 4–6 add coordinated whole-body movement. Direct eye gaze (condition 2) serves as the reference condition for the decomposition, combining genuine social engagement with negligible movement.

EEG was recorded across six conditions, each five minutes in duration, in a configuration that simultaneously recorded the participant and the dance instructor: (1) a resting baseline (eyes open); (2) direct eye gaze, seated face-to-face approximately three feet apart; (3) conversation, focused on planning a shared activity; and three movement conditions in which the pair moved together — (4) teacher-led mirroring, in which the participant reproduced the instructor’s movements; (5) participant-led mirroring, in which the instructor reproduced the participant’s movements; and (6) free improvisation, in which the two moved together, initiating, sustaining, and ending movement in continuous mutual response. The three movement conditions were designed to graduate the interpersonal structure of movement — from synchronous imitation (mirroring) to reciprocal, less-constrained coordination (improvisation) — so that the mirroring-versus-improvisation contrast isolates in-the-moment movement generation from interpersonal coordination. Head-mounted accelerometry was recorded throughout so that movement could be measured rather than assumed.

### EEG acquisition

Dual-brain EEG was recorded simultaneously from the participant and the instructor using two portable 32-channel wet-electrode systems (LiveAmp 32; Brain Products GmbH, Gilching, Germany) at a 500 Hz sampling rate. Electrodes were arranged in the standard 10–20 layout (Fp1, Fp2, F3, F4, F7, F8, Fz, FC1, FC2, FC5, FC6, C3, C4, Cz, T7, T8, FT9, FT10, TP9, TP10, CP1, CP2, CP5, CP6, P3, P4, P7, P8, Pz, O1, O2, Oz). The two amplifiers were hardware-synchronized, sharing a common synchronization-pulse train so that the paired recordings derived from a common time base. Recording was continuous across the whole session, and the six five-minute conditions were segmented offline from the recorded synchronization markers. Tri-axial accelerometry was acquired from dedicated channels on the same LiveAmp amplifiers and therefore shares the EEG time base (see *Accelerometry and movement quantification*).

### EEG preprocessing

Preprocessing was performed on the raw continuous data in EEGLAB (MATLAB) using a pipeline optimized for mobile EEG. Data were band-pass filtered (1–45 Hz; zero-phase Hamming-windowed FIR), screened for noisy channels (removed if flat for more than 5 s, exceeding 4 SD of the remaining channels in noise, or correlating below 0.8 with neighboring channels; removed channels were subsequently interpolated), corrected with Artifact Subspace Reconstruction (0.5 s window, 20 SD threshold), and re-referenced to the full-rank common average. Independent components were derived by Adaptive Mixture Independent Component Analysis (AMICA) and classified with ICLabel. Components for which ICLabel assigned a probability of 0.90 or greater to any single artifact class — muscle, eye, heart, line noise, channel noise, or other — were removed and the remaining components back-projected to the 32 channels, yielding a single artifact-corrected dataset that supports every analysis reported here: channel-level spectral power and parameterization, source-level connectivity, and corticokinematic coherence. This permissive rejection threshold was chosen deliberately, to retain the maximal number of brain components for source reconstruction; a median of 24 components was retained per recording (range 18–30), equal to the effective rank of the resulting data. Following component rejection and re-referencing, each recording was segmented into the six conditions using the recorded synchronization markers, preserving the estimated component weights.

### Effective-rank screening

Because source reconstruction and multivariate connectivity estimation are destabilized by rank deficiency, each artifact-corrected (IC90) recording was screened for rank sufficiency before analysis. The effective rank was computed as the number of singular values exceeding a relative tolerance of 1 × 10⁻⁶, normalized to the largest singular value. Recordings with an effective rank below 15, or containing flat or non-finite channels, were excluded. Although only the multivariate connectivity estimator strictly requires this criterion, it was applied to all neural outcomes so that spectral, connectivity, and corticokinematic-coherence analyses share a common sample; excluded recordings were not retained in the analysis directory and were therefore unavailable to any of them. Of 30 participant recordings screened, three were excluded: two at pre-intervention (effective rank 2 and 11) and one at post-intervention (effective rank 14), leaving 15 pre- and 12 post-intervention recordings. Among retained recordings, effective rank ranged from 18 to 30 (median 24, interquartile range 22–26). Varying the floor to 12 or 18 left the set of participants excluded at both timepoints unchanged; the post-intervention recording with effective rank 14 was retained at a floor of 12 and excluded at 15 and 18. Because the retained brain components form the back-projected basis, each recording’s component count equals its effective rank, so component indices reflect genuine signal rather than null space.

### Spectral power and parameterization

Power spectral density (PSD) was computed from the channel-level time series (IC90 dataset). Each recording was segmented into four-second epochs with 50% overlap. A single epoch length was used for every condition so that frequency resolution, and therefore the low-frequency support that constrains the aperiodic fit, was identical across the conditions being compared; a five-minute condition yields approximately 149 epochs at this setting. For each channel and epoch, the signal was tapered with a Hann window and transformed by Fast Fourier Transform; one-sided power spectra were scaled (doubling non-DC, non-Nyquist bins), normalized by the window’s summed squared coefficients, and averaged across epochs in Welch fashion, which preserves the non-phase-locked oscillatory activity that dominates resting EEG.

Mean spectral power was computed within six non-overlapping bands — delta (1–4 Hz), theta (4–8 Hz), alpha (8–12 Hz), low beta (12–20 Hz), high beta (20–30 Hz), and gamma (30–45 Hz) — and additionally within a broadband beta (12–30 Hz) used only for band-ratio computations and excluded from total-power normalization. Power was expressed in two metrics: absolute power, the log₁₀-transformed band-averaged power (log μV²/Hz), and relative power, the proportion of total power within each of the six non-overlapping bands, where total power is the sum of the six band means across 1–45 Hz. Because movement-related artifact is concentrated at low frequencies, the sensitivity of relative power to the inclusion of delta in this denominator was evaluated directly (**Supplementary Table S1**, **Supplementary Figure S2**).

Region-of-interest (ROI) power was computed by averaging channel power within seven anatomical groupings defined by 10–20 labels: prefrontal (Fp1, Fp2), frontal (F3, F4, F7, F8, Fz, FC1, FC2, FC5, FC6), central (C3, C4, Cz), sensorimotor (C3, C4, Cz, FC1, FC2, FC5, FC6, CP1, CP2, CP5, CP6), temporal (T7, T8, FT9, FT10, TP9, TP10), parietal (P3, P4, P7, P8, Pz, CP1, CP2, CP5, CP6), and occipital (O1, O2, Oz).

The aperiodic component of each channel spectrum was parameterized with SpecParam over 2–44 Hz (peak width limits 1–8 Hz, maximum of six peaks, minimum peak height 0.05, fixed aperiodic mode), yielding an aperiodic exponent and offset together with the power of any periodic peaks. Fitting in knee mode was evaluated as an alternative and rejected: 81% of channel fits returned negative knee parameters, indicating model misspecification over this frequency range, and only 13.5% of fits fell within a physically interpretable range. Model fit statistics for the retained fixed-mode fits are reported by condition in Supplementary Table S2.

Because relative band power is not independent of the aperiodic component, the periodic power of each band was additionally quantified as the mean residual between the observed log-power spectrum and the fitted aperiodic curve, computed over the 2–44 Hz fit range. This measure is continuous and defined for every channel, avoiding the condition-dependent censoring that arises when discrete oscillatory peaks are detected in some recordings but not others. All three metrics are reported for every band and contrast in **Supplementary Table S4**.

### Source reconstruction and intra-brain connectivity

Source-level connectivity was computed from the same IC90 decomposition. Equivalent current dipoles were fit to each retained independent component using EEGLAB’s DIPFIT plugin against a boundary-element head model derived from the Montreal Neurological Institute template brain; components whose scalp topography was bilaterally symmetric were re-fit as two symmetric dipoles. A lead field was computed from the Colin27 template head model included in EEGLAB. Sensor-level data were projected to source space using a linearly constrained minimum-variance (LCMV) beamformer, downsampled to 100 Hz, and parcellated into the 68 cortical regions of the Desikan–Killiany atlas, as implemented in the ROIconnect plugin(Pellegrini et al., 2023). Two source reconstructions were computed from the same leadfield. The multivariate interaction measure and absolute coherence were derived from a three-component-per-region decomposition (nPCA = 3), which the multivariate estimator requires; the orthogonalized amplitude envelope correlation was derived from a single dominant component per region (nPCA = 1), so that each region contributed one time series to the envelope correlation. Both used identical LCMV settings (regularization 0.05, resampled to 100 Hz, Desikan–Killiany atlas).

To separate the phase and amplitude contributions to intra-brain coupling, three connectivity measures were estimated from this common source reconstruction. The primary measure was the multivariate interaction measure (MIM), a multivariate generalization of the imaginary part of coherency that is insensitive to zero-lag (volume-conduction) artifact; MIM was computed from the three dominant principal components per region via a multivariate autoregressive model (model order = 20)(Ewald et al., 2012). To isolate amplitude coupling, orthogonalized amplitude-envelope correlation (AEC) was computed from the single dominant source signal per region: signals were band-pass filtered, Hilbert-transformed, and pairwise-orthogonalized before their amplitude envelopes were correlated, orthogonalization being required within a single brain, where amplitude-envelope correlation is otherwise inflated by zero-lag leakage. As a combined phase-and-amplitude reference, absolute coherence (aCOH) was computed from the same three-component source model; because aCOH retains the zero-lag component, it remains susceptible to volume-conduction leakage and is therefore reported only alongside MIM rather than as a standalone result. Each measure was estimated across the spectrum and averaged within five frequency bands — theta (4–8 Hz), alpha (8–12 Hz), low beta (12–20 Hz), high beta (20–30 Hz), and gamma (30–45 Hz) — yielding a 68 × 68 region-by-region matrix per measure, participant, condition, and band. Source connectivity was computed on data resampled to 100 Hz, so the upper edge of the gamma band approaches the Nyquist limit. To assess whether this affected the gamma estimates, connectivity was recomputed at 200 Hz in a subset of 72 recordings. Phase-based measures were unchanged in absolute terms in both the gamma and high-beta bands (ratios 0.99–1.01), and although amplitude-envelope correlation was uniformly lower at the higher rate, it was reduced by a comparable factor in both bands (0.76 and 0.80) with the rank ordering of recordings preserved (Spearman ρ ≥ 0.99), indicating a property of the envelope estimator rather than a band-specific consequence of anti-alias filtering. Recomputing the condition contrasts at both rates changed the estimated effect sizes by a median of 0.03 (Hedges’ g), with differences distributed symmetrically about zero and largest for the leakage-contaminated coherence measure rather than for the rescaled amplitude measure. Gamma connectivity estimates were therefore not materially affected by the choice of sampling rate.

Each matrix was summarized as a whole-brain mean over all region pairs and, for a network-resolved description, aggregated to the seven canonical Yeo networks (visual, somatomotor, dorsal attention, ventral attention/salience, limbic, frontoparietal control, and default mode). Each Desikan–Killiany region was assigned to its dominant Yeo network using a fixed, published spatial-overlap mapping; network-pair connectivity was computed as the mean over all region pairs spanning the two networks, with within-network values excluding the diagonal, yielding a 7 × 7 network matrix (7 within-network and 21 between-network values) per participant, condition, and band. For each interactive condition, connectivity change was computed as the signed difference from the resting baseline, retaining participants with paired data.

### Accelerometry and movement quantification

Triaxial accelerometry was acquired on dedicated channels of the same LiveAmp amplifiers as the EEG and segmented with the same condition markers, so that movement and neural signals share a common clock. Accelerometry was extracted before EEG filtering and is therefore unfiltered raw data. For each condition, the gravitational component was removed using a fourth-order zero-phase Butterworth high-pass filter (0.3 Hz), the three axes were combined into a single dynamic-acceleration magnitude, and the series was decimated to 50 Hz (Reis et al., 2014). From this magnitude series, we derived (i) movement vigor, the mean dynamic-acceleration magnitude, indexing movement amount; and (ii) movement complexity, quantified as multiscale sample entropy, a complexity index, and spectral entropy, indexing the temporal richness of movement. These indices characterize each individual’s movement; the mirroring and improvisation conditions manipulate its interpersonal structure.

### Corticokinematic coherence

To provide an independent test of whether cortical activity tracked movement — separate from the vigor covariate — corticokinematic coherence (CKC) was computed between each participant’s neural signal and their own head-mounted accelerometer at the fundamental frequency of movement (f0). Both signals were resampled to a common 100 Hz. The movement fundamental was estimated per recording as the peak of the acceleration power spectrum within 0.4–3 Hz, and magnitude-squared coherence was computed by multitaper estimation over 8-s segments with 50% overlap, using three DPSS tapers with a time–bandwidth product of 2. This yields a 0.25 Hz half-bandwidth, narrow enough to separate f0 from its first harmonic at the slow tempi characteristic of this material; coherence was read at f0 and 2·f0 as the maximum within ±0.15 Hz. Recordings shorter than 30 s were rejected.

Statistical reliability was assessed against a within-participant null constructed by shuffling the correspondence between accelerometric and neural segments before estimating coherence, which preserves the spectral content of both signals while destroying their temporal alignment; twenty such draws were taken per recording and condition, and the observed CKC was expressed relative to their mean. To test whether this tracking was somatotopically specific, CKC was compared between sensorimotor channels (FC1, FC2, C3, Cz, C4, CP1, CP2) and the remaining channels; a proprioceptive account predicts a sensorimotor concentration, whereas a diffuse distribution argues against one.

### Behavioral measures

Before and after the intervention, all participants completed a battery of self-report questionnaires targeting empathy (Toronto Empathy Questionnaire, TEQ; Multidimensional Emotional Empathy Scale, MEES), interpersonal functioning (Functional Idiographic Assessment Template Questionnaire, FIAT-Q), social connectedness and belonging (Social Connectedness Scale–Revised, SCS-R; Inclusion of Community in Self Scale, ICS), mindfulness (Five Facet Mindfulness Questionnaire, FFMQ), mood (Profile of Mood States, POMS), perceived stress (Perceived Stress Scale, PSS), anxiety (Beck Anxiety Inventory, BAI), and depression (Beck Depression Inventory, BDI). Attentional control was assessed with a Stroop task. All measures were selected for established validation in adults aged 18 and older. Because the trial was a randomized comparison, group assignment (dance vs. control) was the primary between-subjects factor for behavioral outcomes; analyses used raw summed scores and within-subject pre-to-post change so that each participant served as their own baseline. Full item content and psychometric properties are given in the Supplementary Methods.

#### AI-generated content declaration

During the preparation of this work the authors used Claude in order to identify proper statistical analysis tools and create analysis code; and Undermind in order to conduct a literature review. Figure Labs was used to generate Figure 1. Various AI tools (e.g., ChatGPT, Claude) were utilized for editing written content. After using these tools, the authors reviewed and edited the content as needed and take full responsibility for the content of the published article.

### Statistical analysis

#### Overview

As a pilot, inference was framed around effect-size estimation rather than null-hypothesis significance testing; p-values are reported for transparency but are interpreted alongside effect magnitudes and the consistency of effects across conditions and measures. Effect sizes are reported as within-participant Hedges’ g with 10,000-sample bootstrap 95% confidence intervals (rank-biserial correlations for behavioral Wilcoxon tests) and Spearman ρ for associations. Because the two intervention arms were small (dance n = 7 and control n = 4 with usable post-intervention EEG), group × time (intervention) contrasts were treated as secondary and reported descriptively; the primary neural questions were within-subject — how interactive conditions differ from resting baseline — for which the repeated-measures design provides substantially greater statistical efficiency than the between-group comparison. False-discovery-rate (FDR) control was applied within each test family (within scale for behavioral outcomes; within frequency band for spectral and connectivity outcomes).

This was a pilot study, and the sample size was determined by feasibility rather than by a power calculation: the eighteen participants recruited represent the number that could be enrolled and scanned within the funding period. The within-subject decomposition, which carries the primary inference, contributes 15 participants across six conditions and is powered to detect within-participant effects of approximately d = 0.78 at 80% power (two-tailed, α = .05); effects of that magnitude or larger were observed for the aperiodic, alpha, and amplitude-connectivity contrasts. The intervention arms (7 and 7 at randomization) are underpowered for between-group inference and are reported as effect-size estimates with confidence intervals rather than as hypothesis tests.

#### Movement analysis

Condition effects on movement vigor and complexity were tested as within-participant changes from the resting baseline (Hedges’ g), pooled across arms and time points, and using all available participant-sessions. The synchronous-versus-dynamic structure of the design was tested with a planned contrast of the mirroring conditions (teacher- and participant-led) against free improvisation.

#### Spectral and connectivity condition effects

Whether interactive conditions shifted cortical state relative to rest was tested with linear mixed-effects models fit to each neural outcome, with condition as a fixed effect (resting baseline as the reference level), session (pre/post) as a covariate, and a random participant intercept; the omnibus condition effect was assessed by likelihood-ratio test against a reduced (condition-free) model, and each interactive condition was contrasted against baseline. A planned contrast compared the verbal condition (conversation) against the non-verbal interactive conditions (eye gaze, mirroring, and improvisation) to test whether any state shift was specific to speech, and a second planned contrast compared the mirroring conditions against improvisation. Because it is well powered relative to the group comparison, this within-subject model — rather than a group × time contrast — carried the primary neural inference; the group was added only for the exploratory intervention question and is reported descriptively.

Because the six conditions were presented in a fixed order, condition and time-in-session are perfectly confounded and cannot be separated by covariate adjustment. To assess whether a simple time-on-task trend could account for the condition profiles, each outcome was additionally fit with condition replaced by a linear term for condition position (1–6), and the two models were compared by *F* test on participant-centered values. The same comparison was applied to the instructor’s recordings, which share the condition sequence but differ in familiarity with the paradigm. To separate condition from time in session more directly, each five-minute recording was additionally divided into first and second halves and the spectral measures recomputed for each half using the same 4-s epoching. Continuous drift predicts that a measure changes within a condition at a rate comparable to its change across conditions; a condition effect predicts that it steps at boundaries and is comparatively flat within them. Within-condition change (second half minus first) and between-condition change (last half of one condition against the first half of the next) were therefore compared on a common per-minute scale, and condition was tested against a linear index of position in the session expressed in half-condition units (1–12).

#### Movement artifact control

Because coordinated movement can itself generate correlated cortical signals, every neural effect was tested against concurrent movement. For each outcome, the within-participant change from baseline was correlated with the concurrent change in accelerometric vigor (Spearman ρ) and re-estimated with vigor entered as a covariate in the mixed model; movement was treated as a candidate confound where the vigor association was significant or adjustment materially attenuated the effect. The vigor covariate tests dependence on movement amount; the low-movement eye-gaze condition provides a muscle-minimal probe, and corticokinematic coherence provides an independent test of movement-locked cortical activity.

#### Behavioral analysis

Pre-to-post change was examined within and between groups. Between-group comparisons of change adjusted for baseline using analysis of covariance (ANCOVA), given the non-trivial baseline imbalance expected under small-sample randomization; outcomes with large baseline differences are interpreted with corresponding caution. Effect sizes are the differences between the dance and control groups (Hedges’ g) and the partial eta-squared from the ANCOVA. Within-group change was tested with Wilcoxon signed-rank tests and the matched-pairs rank-biserial correlation. FDR correction was applied within each instrument’s family of subscales.

#### Brain-behavioral analysis

Associations between neural change and behavioral change were examined as exploratory, hypothesis-generating effect-size estimates. Because the subsample with both usable EEG and complete pre-to-post behavior was small (n ≈ 10–13) and because both arms received a socially interactive program, associations were estimated across all participants rather than within arm, using Pearson and Spearman correlations. These analyses were not corrected for multiple comparisons in a confirmatory sense and are reported for transparency; given the sample size, correlations are interpreted as effect-size estimates rather than reliable findings.

#### Software and data availability

Analyses used MATLAB (EEGLAB with the DIPFIT, ICLabel, and ROIconnect plugins) and Python (MNE-Python; specparam/FOOOF for spectral parameterization; statsmodels for mixed-effects models). Analysis code and de-identified derived data required to reproduce all figures and statistics are available at a GitHub repository (https://github.com/embodiedbrainlab/Dance-on-the-Brain); owing to the small, de-identified sample, raw EEG and individual-level behavioral data are available from the corresponding author upon request.

## RESULTS

### Participants

Of 18 adults who consented, four withdrew after the pre-intervention assessment and 14 were randomized to improvisational dance training (n = 7) or a dance-themed movie-watching control (n = 7). Because pre-intervention assessment preceded randomization, participants who subsequently withdrew still contributed a pre-intervention session; and because the primary analyses concern within-session state changes across conditions rather than the intervention contrast, they draw on all completed sessions rather than on intervention completers. Dual-brain EEG was recorded at 17 pre- and 13 post-intervention sessions. Effective-rank screening excluded three participant recordings (DANC pre, ranks 2 and 11; one post, rank 14), leaving 15 pre- and 12 post-intervention sessions for all spectral and connectivity analyses, of which 11 were paired (7 dance, 4 control). One post-intervention session lacked the resting-baseline condition owing to a recording failure and contributed to the five remaining conditions only. Whole-head accelerometry, which is not subject to the rank criterion, was available for all 17 pre- and 13 post-intervention sessions. Thirteen participants completed both behavioral batteries (6 dance, 7 control); one dance participant completed post-intervention EEG but not the questionnaires, and one control participant completed the intervention and questionnaires but never underwent EEG (**Supplementary Table 3**). Participants self-reported both sex and gender identity; both are reported because they diverged in this sample. Of 18 participants, 16 (88.9%) reported female sex and 2 (11.1%) male sex; 15 (83.3%) identified as women, one as a man, one as agender, and one as genderfluid. Racial identification was White (8), Asian (5), Multiracial (2), Black or African American (1), and other (2); 16 (88.9%) reported non-Hispanic ethnicity. Socioeconomic characteristics were not collected. No analyses were stratified by these characteristics, as the sample was too small to support subgroup comparison.

### A graded decomposition of dyadic dance

The six conditions were designed to isolate the components of coordinated social movement. Relative to a solitary resting baseline, direct eye gaze introduced nonverbal social attention while holding the body nearly still; conversation added verbal engagement, again with minimal movement; and the three movement conditions added whole-body coordinated movement, graduating its interpersonal structure from teacher- and participant-led mirroring to free improvisation. Head-mounted accelerometry confirmed the intended movement gradient (**Figure 2**): vigor was at floor during baseline and eye gaze (M = 7.3 mg each; eye gaze vs. baseline g = −0.02, ns), rose modestly during conversation (M = 29.5 mg), and increased sharply across the movement conditions (follow M = 126.5, lead M = 149.0, improvisation M = 212.6 mg; all p < .001), with improvisation both more vigorous (g = +1.15) and more temporally complex (g = +0.67) than mirroring. Because eye gaze combines genuine social engagement with negligible movement, it provides the natural reference state for the analyses that follow: each subsequent component of dance is measured as a change relative to nonverbal social attention rather than to solitary rest. We report four participant-side measures: the aperiodic exponent and relative band power at the channel level, and, at the source level, amplitude-envelope correlation (AEC) and the leakage-robust multivariate interaction measure (MIM), which index amplitude and phase coupling respectively.

**Figure 2.**
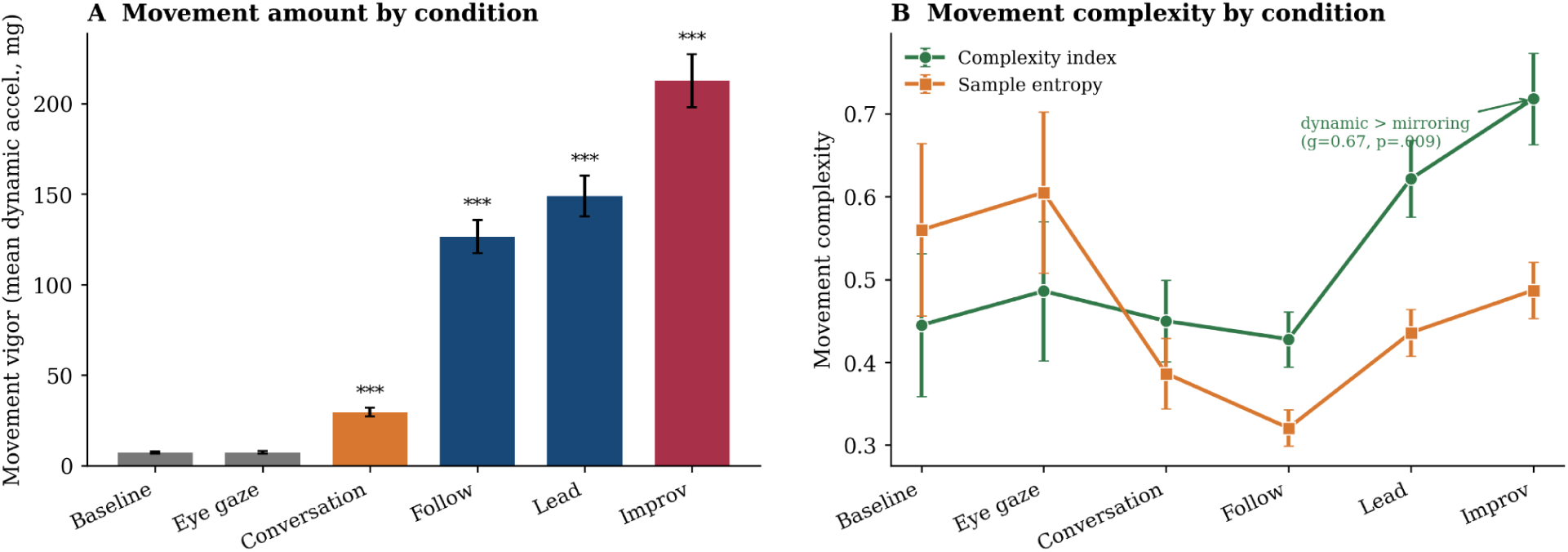
Movement outcomes confirm the intended gradient. (A) Movement vigor (mean dynamic acceleration) by condition, at floor during baseline and eye gaze and rising through the movement conditions to a peak at improvisation (*** p < .001 vs. baseline). (B) Movement complexity (complexity index and sample entropy): free improvisation is more temporally complex than the mirroring conditions. Because eye gaze combines genuine social engagement with negligible movement, it serves as the reference state for the decomposition in Figures 3–4. Bars/points show mean ± SEM; n = 30 participant-sessions.

Referenced this way, the neural changes partition cleanly across the components of the interaction (**Figures 3 and 4**). Nonverbal social attention accounted for most of the shift in cortical state; verbal engagement added a large amplitude-connectivity effect; coordinated movement added fast-band power and a network reorganization; and generative improvisation, examined as an exploratory contrast, reorganized frontoparietal coupling. For clarity, we assess each component in turn (**Table 1**, **Figure 3**), with the incremental contribution of each tier summarized in **Figure 4**.

**Figure 3.**
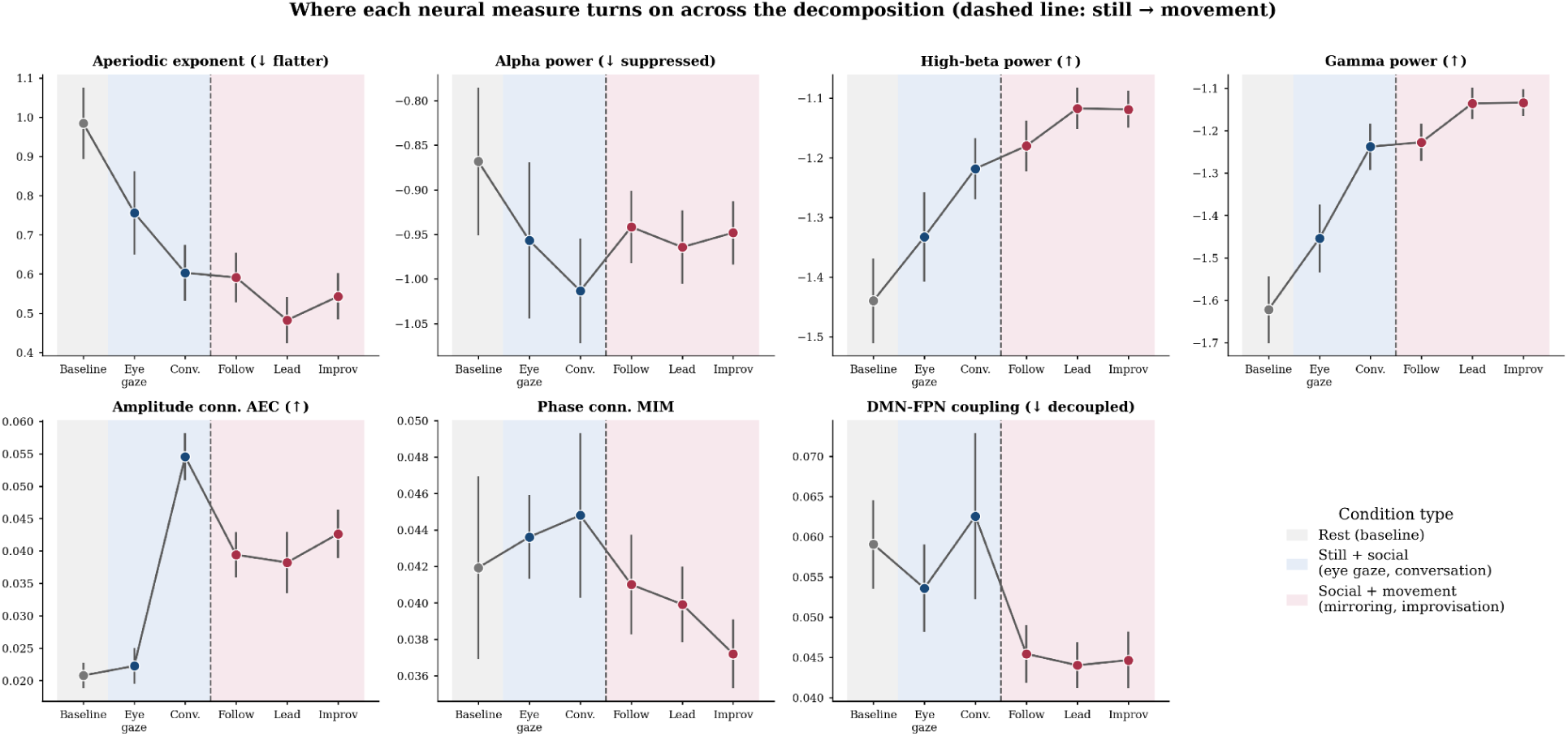
Neural measures across the graded conditions. Each panel shows one participant-side neural measure (channel-level aperiodic exponent; relative alpha, high-beta, and gamma power; whole-brain high-frequency amplitude connectivity [AEC] and phase connectivity [MIM]; and theta/alpha default-mode–frontoparietal [DMN–FPN] phase coupling) across the six conditions, ordered from rest through the still-social conditions to the movement conditions. Background shading denotes condition type (rest; still + social; social + movement), and the dashed line marks the still → movement transition. The aperiodic, alpha/mu, and fast-band spectral changes emerge during the still social conditions, whereas amplitude connectivity and the DMN–FPN decoupling emerge with verbal engagement and movement. Points show means ± SEM (pre-intervention; n = 15).

**Figure 4.**
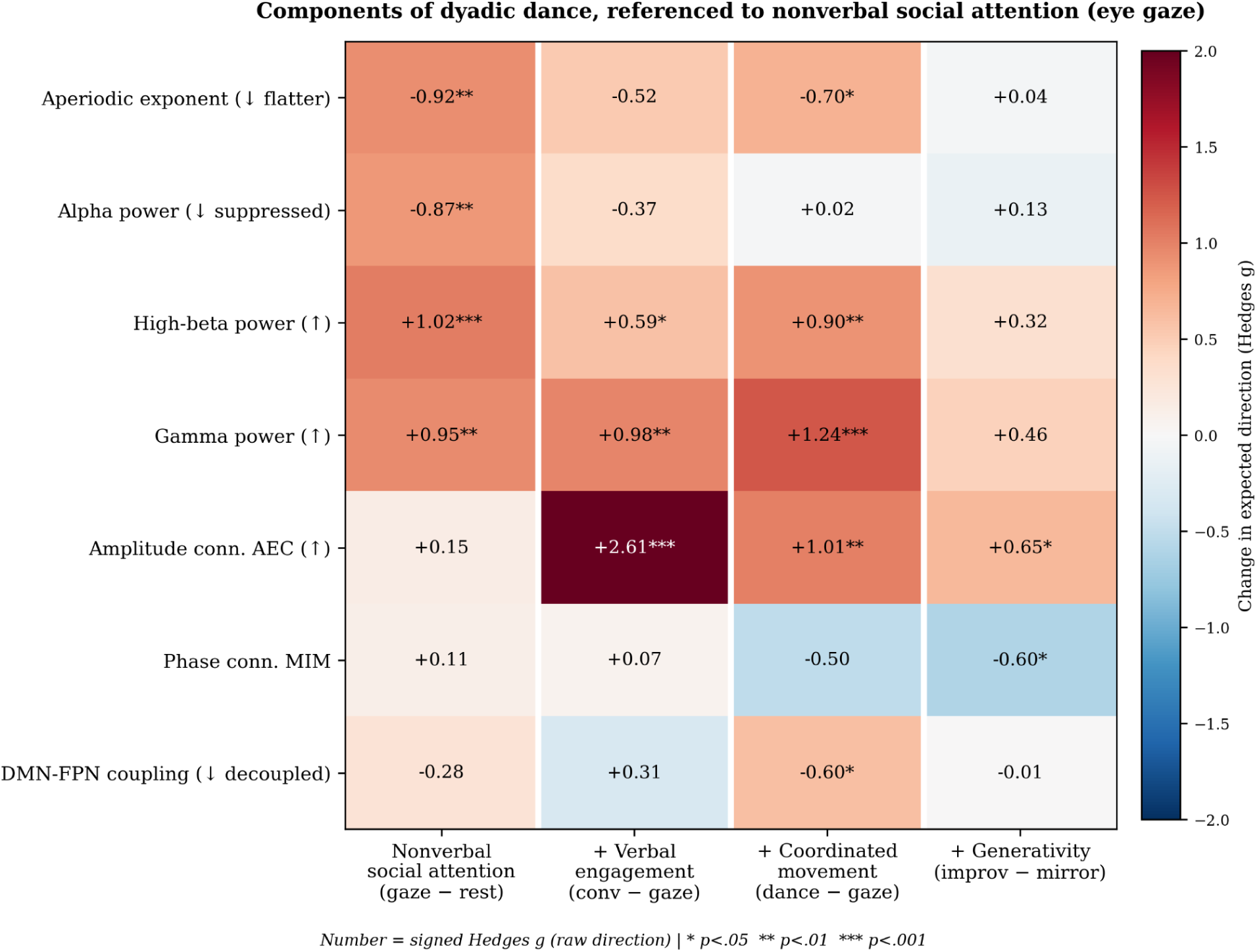
Components of dyadic dance. Each neural measure (rows) is decomposed into the incremental contribution of nonverbal social attention (eye gaze − baseline), verbal engagement (conversation − eye gaze), coordinated movement (movement conditions − eye gaze), and generative improvisation (improvisation − mirroring; columns). Cell color encodes the change in the expected direction for each measure (red = state change; blue = opposite), and the printed number is the signed within-participant Hedges’ g (raw direction). Nonverbal social attention accounts for most of the spectral state change; verbal engagement adds amplitude connectivity; coordinated movement adds fast-band power and DMN–FPN decoupling; and generative improvisation reorganizes frontoparietal coupling (exploratory; see Figure 6). Arrows in the row labels indicate the direction of the expected effect. * p < .05, ** p < .01, *** p < .001; pre-intervention, n = 15.

### Non-verbal social attention establishes the engaged brain state

Simply attending to a partner, without speaking or moving, produced most of the cortical state change observed anywhere in the paradigm. Relative to solitary baseline, eye gaze flattened the aperiodic exponent (g = −0.97, p = .001), suppressed alpha power (relative g = −0.87, p = .003), and increased relative high-beta (g = +1.02, p < .001) and gamma power (g = +0.95, p = .002). Because relative power is not independent of the aperiodic fit — relative gamma and high beta correlate with the exponent at rho = −0.95 and −0.94 across condition cells — we additionally quantified the periodic component of each band as the residual power above the fitted aperiodic curve (**Supplementary Table 4)**. The alpha reduction was confirmed as an oscillatory change (periodic alpha g = −0.90, p = .002) and was distributed broadly across the scalp rather than being focal to sensorimotor cortex (all seven regions p < .05 after correction; range −0.57 to −1.09). The high-beta increase did not survive aperiodic adjustment (periodic g = +0.28, p = .27), indicating that it reflects the spectral tilt rather than a change in oscillatory activity, whereas the gamma increase was partly oscillatory (periodic g = +0.57, p = .01). In contrast, neither amplitude connectivity (AEC g = +0.15, p = .55) nor phase connectivity (MIM g = +0.11, p = .65) changed with nonverbal social attention alone. Thus the aperiodic and alpha signatures of engagement were largely established the moment two people attended to one another, whereas coordinated inter-regional connectivity was not yet engaged.

### Verbal engagement adds amplitude connectivity

Adding verbal engagement, while movement remained minimal, produced a further, specific change: a large increase in high-frequency amplitude connectivity relative to eye gaze (AEC g = +2.61, p < .001; **Supplementary Figure S3**), accompanied by a further rise in gamma power (g = +0.91, p = .002) and high-beta power (g = +0.51, p = .056). The aperiodic exponent flattened marginally further (g = −0.48, p = .070), and alpha suppression did not deepen reliably (g = −0.47, p = .073). Conversation is thus distinguished from silent social attention chiefly by the emergence of amplitude-based coupling. As shown below, this amplitude-connectivity effect co-varies with movement and is the most susceptible of the connectivity measures to a speech-related myogenic contribution, so it is interpreted with corresponding caution.

### Coordinated movement adds fast-band power and network reorganization

Coordinated whole-body movement, relative to nonverbal social attention, added two things. First, it deepened the spectral shift already established by social attention: the aperiodic exponent flattened further (g = −0.61, p = .026), and fast-band power increased — gamma (g = +1.13, p < .001) and high-beta (g = +0.78, p = .007) — as did amplitude connectivity (AEC g = +1.01, p = .001). Second, and in contrast to the still conditions, it reorganized phase-based network coupling: theta/alpha coupling between the default-mode and frontoparietal-control networks decreased during movement relative to eye gaze (DMN–FPN g = −0.59, p = .029; **Supplementary Figure S4**), and whole-brain high-frequency phase coupling trended lower (MIM g = −0.50, p = .059). This contrast was specified a priori on the basis of the task-positive/task-negative account rather than selected from the full set of network pairs. Across all 28 pairs, the movement-related decrease was broadly distributed and centred on the frontoparietal-control network, which appeared in five of the eight largest decreases (**Supplementary Figure S4**); no individual pair survived correction across the full set. Alpha suppression, already saturated by social attention, did not change further with movement (g = −0.07, p = .77), although this null is the one spectral result sensitive to the treatment of delta power in the relative-power denominator (Supplementary Table S1). Coordinated movement therefore contributes a fast-band, amplitude-weighted intensification together with a broad low-frequency withdrawal of frontoparietal-control connectivity — the network signature of a more strongly externally directed state.

### Distinguishing which effects are driven by movement, and which by engagement

Because three of the four tiers involve movement, we asked which effects depend on movement amount and which reflect social engagement independent of it, using concurrent movement vigor as a covariate, the near-still eye-gaze condition as a muscle-minimal probe, and corticokinematic coherence (CKC) as an independent index of movement-locked cortical activity. The effects separated consistently. The aperiodic flattening and alpha suppression were movement-independent: both were already maximal during eye gaze, neither scaled with vigor across conditions (aperiodic ρ = −0.13, p = .13; alpha ρ = −0.01, p = .67), and the flattening survived vigor adjustment region by region, including over sensorimotor cortex. The theta/alpha DMN–FPN decoupling was likewise unrelated to vigor (ρ = −0.10 to −0.17, ns). In contrast, the fast-band power increases, while surviving vigor adjustment and present during eye gaze, showed a genuine movement-linked component (gamma ρ = +0.28, p < .001, after adjusting for condition), and the amplitude-connectivity (AEC) increase was the most movement-dependent effect: it was absent during eye gaze (g = +0.15, ns), co-varied with vigor (ρ = +0.37, p < .001), and rose most during conversation, consistent with a speech-related contribution (**Figure 5**).

**Figure 5.**
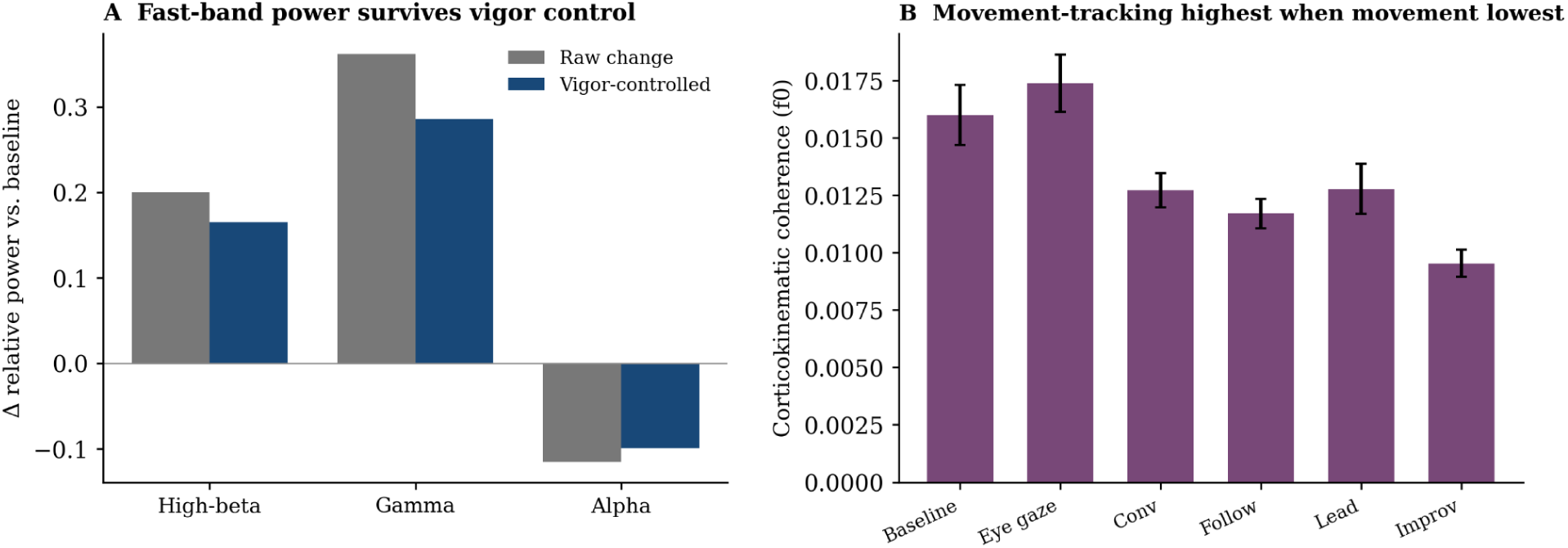
The core effects are separable from movement and muscle. (A) Relative fast-band power change, shown raw and after adjusting for concurrent movement vigor: the gamma and high-beta increases survive vigor adjustment (a significant residual remains), while alpha suppression is vigor-independent. (B) Corticokinematic coherence (coherence between each participant’s cortex and their own accelerometer at the movement fundamental) by condition: cortical tracking of the movement rhythm is highest during the low-movement conditions and lowest during the most vigorous condition (improvisation) — the opposite of a movement-artifact gradient. Together with the presence of the aperiodic, alpha, and DMN–FPN effects during near-still eye gaze, these controls indicate that the social-attention and network components of the decomposition are cortical rather than myogenic, whereas the fast-band and amplitude-connectivity components carry an expected movement-linked contribution. Means ± SEM.

Corticokinematic coherence reinforced this separation. Cortical activity tracked the movement rhythm above chance (mixed-model estimate = 0.017, p < 10⁻¹⁰), but this tracking was not localized to sensorimotor cortex (sensorimotor vs. non-sensorimotor g = −0.02, p = .81) and, counter to a myogenic account, was strongest during the low-movement conditions (eye gaze, baseline) and weakest during the most vigorous condition (improvisation). Brain state tracked movement amount rather than movement complexity throughout: neither the aperiodic flattening nor the fast-band increase was predicted by movement complexity once vigor was accounted for.

Because relative power expresses each band as a proportion of total power, and because movement-related artifacts are concentrated at low frequencies, we tested whether the relative-power results depended on the inclusion of delta in that denominator. Absolute power increased with movement vigor in every band, but the raw association across all condition cells was strongest for gamma (ρ = +0.66) and high beta (ρ = +0.60) and weaker for delta (ρ = +0.45), a gradient more consistent with a myogenic than a low-frequency kinematic contribution; relative delta was unrelated to vigor (ρ = +0.01, p = .86), indicating that delta rose in approximate proportion to total power rather than disproportionately inflating the denominator. Recomputing relative power with delta excluded left the fast-band results essentially unchanged (gamma and high beta shifted by ≤ 0.05 at every contrast) and did not alter the eye-gaze effects, but produced a larger alpha suppression during the movement conditions (g = −0.41 versus −0.07 for movement relative to eye gaze). The full comparison is given in **Supplementary Table S1** and **Supplementary Figure S2**. We retain the conventional full-band denominator throughout, and note that the conclusion that alpha suppression is established by social attention rather than by movement is the one claim that is weakened under the alternative normalization. Because the conditions were presented in a fixed order, we also asked whether a linear time-on-task trend could account for the condition profiles as well as condition itself. For the aperiodic exponent, periodic alpha power, and amplitude connectivity, condition explained significantly more variance than a linear trend (F(4,155) = 8.47, 7.23, and 37.43, respectively, all p < .001); amplitude connectivity was reliably higher during conversation than during teacher-led mirroring (g = +0.77, p = .005), a reversal that a monotone trend cannot produce. For the theta/alpha frontoparietal decoupling and for corticokinematic coherence, a linear trend fitted the data as well as condition (F = 1.37 and 0.94, both ns), so a contribution of time in session to those two gradients cannot be excluded. Dividing each recording into halves separated the two accounts directly. The pooled within-condition change in the aperiodic exponent was negligible (+0.006, g = +0.04, p = .687), whereas the mean change across condition boundaries was 23 times larger per unit time, with the largest steps at the baseline-to-eye-gaze (g = −1.04, p = .001) and eye-gaze-to-conversation (g = −0.78, p = .007) transitions. Condition also explained variance beyond a linear index of session position measured in half-condition units (F(5,173) = 7.96, p < .001; R² increased from 0.36 to 0.48). One condition showed reliable within-condition change: the exponent rose during eye gaze (g = +0.72, p = .011), partially returning toward baseline across the five minutes, a pattern consistent with relaxation of an initially intense state rather than with monotonic drift. As a further check, the same analysis was applied to the instructor, who completed this identical condition sequence on approximately thirty occasions and for whom the paradigm was therefore not novel. The movement-related decrease in frontoparietal coupling was present in the instructor at comparable magnitude (g = −0.55, p = .004), indicating that novelty, practice, or expectation effects do not account for the gradient. As a check on the leakage-robustness of the connectivity measures, we also examined absolute coherence, which includes the zero-lag component and is therefore inflated by volume conduction within a single brain. It increased modestly and non-significantly across the interactive conditions (eye gaze g = +0.50; conversation g = +0.41; movement g = +0.34) and was uncorrelated with both leakage-robust measures across condition cells (AEC ρ = +0.10; MIM ρ = −0.07). That it tracks neither the amplitude nor the phase measure is consistent with its susceptibility to volume conduction, and inferences about coordinated neural activity therefore rest on MIM and AEC. Together, these controls — vigor adjustment, the muscle-minimal eye-gaze probe, corticokinematic coherence, the denominator check, the time-on-task comparison, and the leakage-robustness check — indicate that the social-attention and network components of the decomposition are genuine cortical effects, while the fast-band and amplitude-connectivity components carry a real and expected contribution from movement itself.

### Generative improvisation as an exploratory contrast

Finally, we compared free improvisation with structured mirroring to ask whether generating movement in the moment, rather than reproducing a partner’s movements, carried a distinct signature. Because improvisation was also the most vigorous and most complex condition, this contrast is confounded with movement amount and is reported as exploratory. Relative to mirroring, improvisation shifted the balance of high-frequency connectivity from phase to amplitude: amplitude connectivity increased (gamma AEC g = +0.76, p = .008) while phase connectivity decreased (low-beta MIM g = −0.74; gamma MIM g = −0.76, both p < .01). This reorganization was concentrated on the frontoparietal-control network (**Figure 6**): the amplitude increase was carried by DAN–FPN, somatomotor–DAN, and default-mode connections, with the phase decrease carried by connections into the FPN (limbic–FPN, ventral-attention–FPN, FPN–default-mode, somatomotor–FPN). The pattern is consistent with the greater executive and attentional demands attributed to improvisation, but its dependence on movement limits interpretation: the amplitude-connectivity increase survived adjustment for movement complexity (p = .039) but not for movement vigor (p = .92), indicating that it tracks how much participants moved rather than generative demand alone. We therefore regard the frontoparietal reorganization as a promising but movement-confounded lead for future work in which movement amount is equated across conditions.

**Figure 6.**
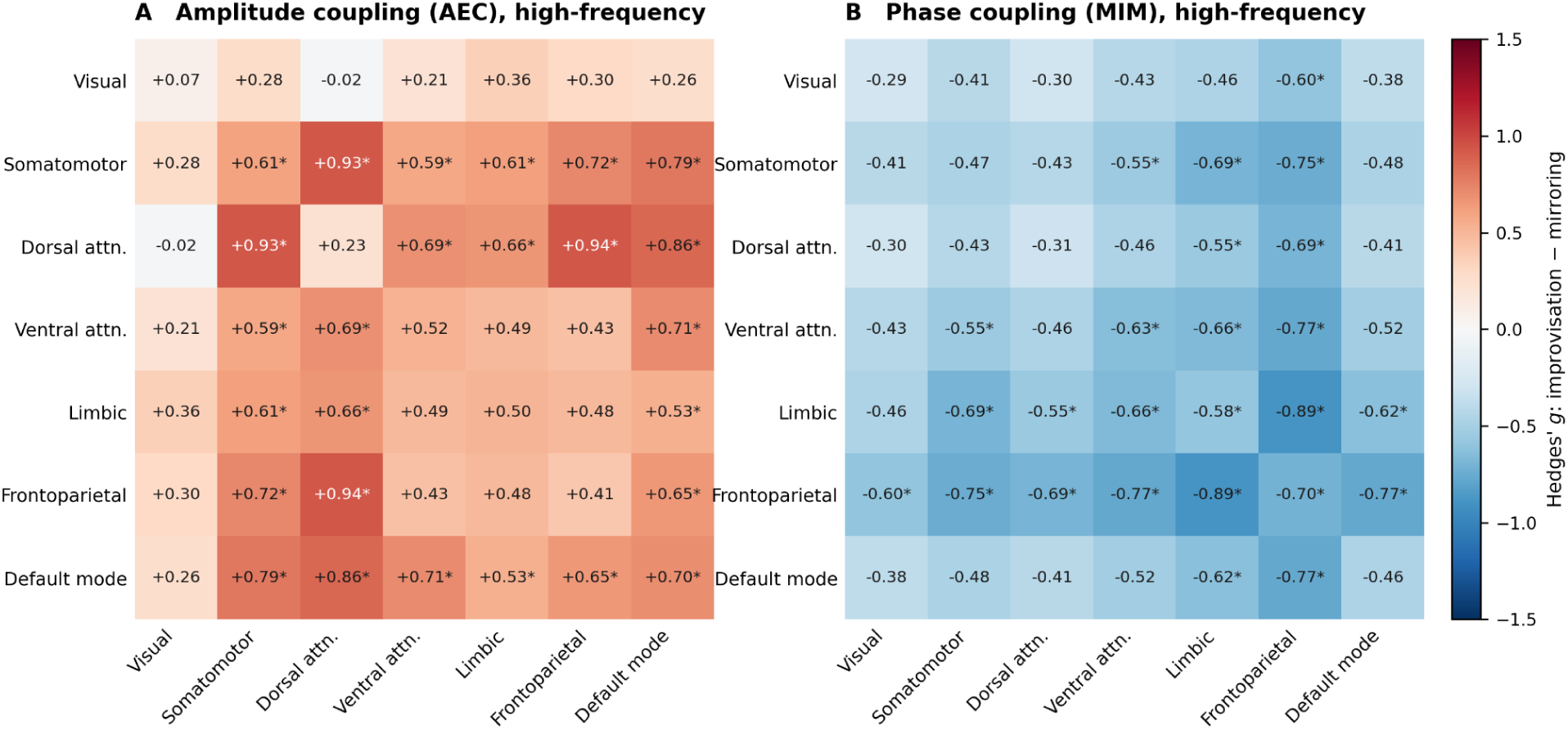
Improvisation versus mirroring. Network-resolved high-frequency (high-beta + gamma) connectivity change for free improvisation relative to structured mirroring, across seven canonical Yeo networks. (Left) Amplitude coupling (AEC) increases during improvisation, concentrated on dorsal-attention–frontoparietal (DAN–FPN), somatomotor–DAN, and default-mode connections. (Right) Phase coupling (MIM) decreases during improvisation, concentrated in connections to the frontoparietal-control network (limbic–FPN, ventral-attention–FPN, FPN–default-mode, somatomotor–FPN). Because improvisation was also the most vigorous condition and the amplitude effect did not survive adjustment for movement vigor, this contrast is reported as exploratory and movement-confounded. Cells show within-participant Hedges’ g (improvisation − mirroring); pre-intervention, n = 15.

### Intervention effects (group x time)

Because participants were randomized to dance versus control, group × time effects were examined. With few participants completing post-intervention EEG per arm (dance n = 7, control n = 4), these contrasts were underpowered and yielded no reliable effects (aperiodic flattening, dance vs. control change p = .53; AEC p = .79); they are reported descriptively and not interpreted as intervention effects.

### Behavioral outcomes

Four weeks of improvisational dance produced medium-to-large effects favoring the dance group on four of five outcomes — stress, empathy for others’ distress, interpersonal conflict, and attentional performance — with no group difference in social connectedness (**Table 2**). After within-scale FDR correction, one effect survived: on the MEES, the dance group increased in responsive-crying empathy while the control group decreased (g = 0.82; ANCOVA p = .006, η²p = 0.54; corrected p = .044). Additional large effects in the hypothesized direction, which did not survive correction, were observed for interpersonal conflict (FIAT-Q g = −1.08, p = .031), perceived stress (PSS g = −0.99, p = .056), and Stroop congruent reaction time (g = −0.91, p = .085). Social connectedness did not differ between groups (g = −0.02).

### Brain-behavioral relationships (exploratory)

To connect neural state changes to behavior, participant-level change indices for aperiodic flattening, gamma power, and alpha power were correlated with pre-to-post changes in behavioral measures. Given the small paired subsample (n ≈ 10), these associations are exploratory effect-size estimates, and none survived FDR correction; the strongest were in the stress domain (gamma-change × stress-change r = +0.63, p = .050; flattening × stress-change r = −0.61, p = .064). They are consistent with the possibility that the magnitude of the interaction-induced state change tracks the degree of behavioral change, but require adequately powered replication.

## DISCUSSION

We used mobile hyperscanning EEG and a graded set of dyadic conditions to decompose the neural signature of coordinated social movement. By referencing each condition to nonverbal social attention — direct eye gaze, which combines genuine social engagement with negligible movement — rather than to solitary rest, we could ask which features of the individual-brain response belong to social attention itself, which to verbal engagement, which to coordinated movement, and which to the generative demands of improvisation. The ordering was unexpected: most of the shift in cortical state was established by nonverbal social attention alone, before any movement occurred.

### Nonverbal social attention establishes the engaged-brain state

Simply attending to another person, without speaking or moving, reproduced most of the cortical state change observed across the paradigm, locating the core of the effect in social attention rather than movement. The flattening of the aperiodic exponent is widely interpreted as a shift of cortical dynamics toward excitation and heightened arousal(Gao et al., 2017), and tracks arousal and neuromodulatory tone more broadly(He, 2014; Lendner et al., 2020). That this flattening appeared during still eye gaze indicates that the arousal it reflects is social rather than motor in origin. Direct mutual gaze is a potent social signal that rapidly engages the social brain and modulates ongoing cognition and autonomic arousal(Senju and Johnson, 2009).

Alpha functions as an inhibitory gating rhythm whose desynchronization releases cortical regions for active processing(Jensen and Mazaheri, 2010; Klimesch, 2012). The reduction observed here was confirmed as a change in oscillatory power rather than a consequence of the aperiodic shift, but was broadly distributed rather than focal to sensorimotor cortex, and is therefore not specific enough to identify with the mu rhythm, which in action-observation paradigms shows a centroparietal decrease that spares occipital cortex(Oberman et al., 2007; Fox et al., 2016). Whether sensorimotor resonance contributes remains open. The accompanying rise in high-frequency power should be read with the aperiodic result rather than as independent evidence: after aperiodic adjustment the high-beta increase did not persist and the gamma increase was roughly halved. What these data establish is that a measurable cortical state accompanies mutual attention in the absence of movement. In the vocabulary of dance/movement therapy, kinesthetic empathy and social attunement describe the embodied sharing of another’s presence(Fischman, 2015); these data ground that idea by showing that attention to a partner alone reorganizes cortical state.

### Verbal engagement adds amplitude-based connectivity

Adding verbal engagement, while the body remained nearly still, was distinguished chiefly by the emergence of high-frequency amplitude connectivity, which had not changed with silent social attention. Conversation recruits a distributed frontotemporal network with particular prominence of gamma-band activity during both production and comprehension in natural dialogue(Park et al., 2015; Cai et al., 2025), so a rise in high-frequency amplitude coupling is plausibly in part a genuine correlate of engaging these systems. Our control analyses nonetheless indicate that this is the most movement- and muscle-susceptible of the connectivity measures, co-varying with vigor and largest precisely when participants were speaking. Speech entails continuous orofacial movement whose myogenic activity projects to frontotemporal electrodes and can inflate broadband high-frequency power and its inter-regional correlation. We therefore interpret this effect cautiously and rely on the phase-based measure, which is not susceptible to this contamination, for inferences about coordinated neural activity.

### Coordinated movement adds fast-band power and network reorganization

Coordinated whole-body movement contributed a further increase in fast-band power and amplitude connectivity, and, in contrast to the still conditions, a reorganization of phase-based network coupling in which theta/alpha coupling involving the frontoparietal-control network decreased, including its coupling with the default-mode network. This recapitulates the task-positive/task-negative organization described for solitary externally directed tasks, in which engagement of attentional systems accompanies disengagement of the default-mode network(Fox et al., 2005), with frontoparietal cortex arbitrating according to task demands(Yin et al., 2022); that the effect fell in theta and alpha aligns with intracranial evidence that this antagonism is carried by low-frequency activity(Hammer et al., 2024). Unlike the fast-band and amplitude effects, this decoupling was independent of movement vigor, indicating that it reflects the coordinative demand of moving with a partner rather than motor load. Joint action requires continuous reciprocal prediction and monitoring(Sebanz et al., 2006), which places a premium on external attention. This contrast was specified a priori from the task-positive/task-negative account rather than selected from the full set of network pairs. Across all 28 pairs the movement-related decrease was broadly distributed and centred on the frontoparietal-control network, which appeared in five of the eight largest decreases (**Figure S4**); no individual pair survived correction across the full set. This gradient is, however, among the measures for which a time-on-task contribution cannot be excluded.

### Generative improvisation and frontoparietal reorganization

Comparing free improvisation with structured mirroring asked whether generating movement in the moment carried a distinct signature. Improvisation shifted the balance of high-frequency connectivity from phase to amplitude, concentrated in the frontoparietal-control network — consistent with the executive and attentional demands attributed to improvisation in musical and motor domains(Limb and Braun, 2008; Beaty, 2015). Improvisation was, however, also the most vigorous and complex condition, and the amplitude-connectivity increase survived adjustment for movement complexity but not for movement vigor. This contrast is therefore confounded with movement amount and reported as exploratory.

### Interpreting the decomposition

Read as a whole, “dancing together” is not a single neural event but a layered one. The components of a real interaction do not contribute equally: nonverbal social attention establishes an aroused, attentive cortical state — the bulk of the spectral change — before any movement occurs. Coordinated movement then adds a reorganization of large-scale connectivity toward an externally directed configuration. Verbal engagement and generative improvisation contribute further, more specific changes. This refines the prediction of the Synchronicity Hypothesis of Dance(Basso et al., 2020): rather than a generalized increase in neural communication, coordinated social movement produces a selective and componential reconfiguration in which the socially engaged state and the coordinative network shift are separable and evoked by different features of the encounter.

That social attention accounts for most of the state change suggests the engaged state is not contingent on vigorous movement but is available whenever two people attend and coordinate through gaze and attunement. Practices built on mirroring, postural coupling, and shared attention can be understood as ways of inducing it, and the componential structure offers a vocabulary for asking which element of a given practice is doing the work.

### Socio-emotional outcomes

Four weeks of improvisational dance produced medium-to-large effects on four of five socio-emotional outcomes, most reliably an increase in empathic responsiveness to others’ distress, with convergent effects on perceived stress, interpersonal conflict, and attentional performance, and no group difference in social connectedness. Because the sample was small and the intervention arms smaller still, these are reported as effect-size estimates rather than definitive intervention effects. Their direction is coherent with the neural findings: a practice that repeatedly engages the systems supporting attention to another person might be expected to strengthen empathic capacities. The exploratory brain–behavior associations require adequately powered replication.

### Limitations

The sample was small and the intervention arms smaller still, so group-by-time analyses were underpowered and are reported descriptively; the decomposition rests on better-powered within-subject comparisons, but replication in a larger and more diverse sample is needed, particularly given a predominantly female, non-Hispanic sample from a single region. The six conditions were presented in a fixed order, so condition and time-in-session are perfectly confounded and cannot be separated by covariate adjustment. Condition explained the data better than a linear time-on-task trend for the aperiodic, alpha, and amplitude-connectivity effects, and a split-half analysis showed that the aperiodic exponent steps at condition boundaries rather than drifting within them, changing 23 times faster across boundaries than within conditions. The frontoparietal and corticokinematic gradients were not distinguished from a linear trend, though the frontoparietal decrease was present in the instructor, for whom the sequence was highly familiar, so novelty and practice effects are unlikely to explain it. The verbal-engagement and generativity contrasts are confounded with speech-related movement and movement amount respectively. Brain–behavior associations were exploratory and uncorrected in a small paired subsample. Finally, source estimates from a 32-channel montage have limited spatial resolution, so network-level localization should be read as coarse-grained.

### Conclusions

The individual-brain signature of coordinated social movement is layered and componential. Nonverbal social attention alone establishes most of the shift in cortical state toward excitation and attentional engagement; coordinated movement adds a reorganization of large-scale connectivity toward an externally directed configuration; and verbal engagement and generative improvisation contribute further, more specific effects. This favors the selective reconfiguration anticipated by the Synchronicity Hypothesis of Dance over a generalized increase in neural communication, and suggests that the socially engaged state that movement-based practices aim to produce is evoked by attention and coordination themselves, and only intensified by vigorous movement. The aperiodic exponent, alpha-band oscillatory power, and low-frequency frontoparietal coupling emerge as candidate markers for adequately powered trials.

## Supporting information

Manuscript Tables

## Acknowledgments

The authors express their gratitude to Professor Rachel Rugh for guiding the movement exercises and generously allowing her brain to be studied.

## Ethics approval statement

The protocol was approved by the Virginia Tech Institutional Review Board (approval #21-798, approved 6/20/23), and all participants provided written informed consent in accordance with the Declaration of Helsinki.

## CRedit authorship contribution statement

Noor Tasnim: Data curation, Software, Methodology, Formal analysis, Writing – review and editing. Rachel M. DeLauder: Investigation, Formal analysis, Writing – review and editing. Mackenzie Aychman: Investigation, Data curation, Writing – review and editing. Mary Gahagan: Investigation, Writing – review and editing. Jessica Purevtugs: Investigation, Writing – review and editing. Jesse Newpol: Investigation, Writing – review and editing. Ryan Frank: Investigation, Writing – review and editing. Grace Grizzell: Investigation, Writing – review and editing. Grace Nobriga: Investigation, Writing – review and editing. Sarah Rose: Investigation, Writing – review and editing. Julia C. Basso: Conceptualization, Methodology, Formal analysis, Investigation, Resources, Supervision, Project administration, Funding acquisition, Visualization, Writing – original draft, Writing – review and editing.

## Data and code availability

Analysis code supporting the findings of this study is available at https://github.com/embodiedbrainlab/Dance-on-the-Brain. Raw electroencephalographic recordings are not publicly deposited because participants did not consent to public release of identifiable neural and video data; de-identified recordings and derived data are available from the corresponding author on reasonable request, subject to a data use agreement.

