## Supplementary Tables for "Social attention and coordinated movement produce dissociable spectral and network changes during dyadic improvisational dance"

**Table S1. Sensitivity of relative-power effect sizes to the inclusion of delta in the total-power denominator.**

| Band | Contrast | g [95% CI], delta included | q | g [95% CI], delta excluded | q |
|---|---|---|---|---|---|
| Delta | Eye gaze – baseline | -0.31 [-0.80, +0.16] | 0.229 | -0.29 [-0.83, +0.18] | 0.298 |
| Delta | Conversation – eye gaze | +0.36 [-0.10, +0.96] | 0.237 | +0.21 [-0.27, +0.84] | 0.576 |
| Delta | Movement – eye gaze | -0.07 [-0.64, +0.42] | 0.772 | -0.22 [-0.76, +0.28] | 0.448 |
| Delta | Improvisation – mirroring | -0.53 [-1.21, -0.07] | 0.105 | -0.61 [-1.27, -0.16] | 0.079 |
| Theta | Eye gaze – baseline | -0.32 [-0.94, +0.14] | 0.229 | -0.34 [-0.90, +0.12] | 0.271 |
| Theta | Conversation – eye gaze | -0.01 [-0.78, +0.41] | 0.964 | +0.03 [-0.67, +0.45] | 0.894 |
| Theta | Movement – eye gaze | +0.42 [-0.03, +0.88] | 0.166 | +0.06 [-0.49, +0.53] | 0.795 |
| Theta | Improvisation – mirroring | +0.78 [+0.32, +1.53] | 0.039 | +0.21 [-0.26, +0.93] | 0.593 |
| Alpha | Eye gaze – baseline | -0.87 [-1.69, -0.51] | 0.006 | -0.98 [-1.87, -0.65] | 0.004 |
| Alpha | Conversation – eye gaze | -0.47 [-1.07, -0.03] | 0.146 | -0.77 [-1.26, -0.45] | 0.014 |
| Alpha | Movement – eye gaze | -0.07 [-0.63, +0.43] | 0.772 | -0.41 [-0.94, +0.07] | 0.195 |
| Alpha | Improvisation – mirroring | +0.23 [-0.29, +0.69] | 0.358 | -0.73 [-1.41, -0.28] | 0.060 |
| Low Beta | Eye gaze – baseline | +0.31 [-0.18, +0.83] | 0.229 | +0.20 [-0.29, +0.80] | 0.434 |
| Low Beta | Conversation – eye gaze | +0.07 [-0.58, +0.48] | 0.927 | +0.18 [-0.32, +0.72] | 0.576 |

| | | | | | |
|---|---|---|---|---|---|
| Low Beta | Movement – eye gaze | +0.42 [+0.04, +0.85] | 0.166 | +0.39 [-0.06, +0.95] | 0.195 |
| Low Beta | Improvisation – mirroring | +0.48 [+0.04, +0.98] | 0.106 | +0.06 [-0.43, +0.66] | 0.814 |
| High Beta | Eye gaze – baseline | +1.02 [+0.54, +1.88] | 0.005 | +1.00 [+0.52, +1.88] | 0.004 |
| High Beta | Conversation – eye gaze | +0.51 [+0.08, +1.01] | 0.146 | +0.80 [+0.32, +1.57] | 0.014 |
| High Beta | Movement – eye gaze | +0.78 [+0.40, +1.65] | 0.019 | +0.83 [+0.31, +1.75] | 0.013 |
| High Beta | Improvisation – mirroring | +0.38 [-0.09, +0.89] | 0.165 | +0.11 [-0.43, +0.62] | 0.795 |
| Gamma | Eye gaze – baseline | +0.95 [+0.42, +1.95] | 0.005 | +0.92 [+0.40, +1.87] | 0.004 |
| Gamma | Conversation – eye gaze | +0.91 [+0.51, +1.63] | 0.014 | +1.16 [+0.71, +2.02] | 0.002 |
| Gamma | Movement – eye gaze | +1.13 [+0.78, +2.20] | 0.002 | +1.08 [+0.66, +1.77] | 0.003 |
| Gamma | Improvisation – mirroring | +0.52 [+0.07, +1.06] | 0.105 | +0.35 [-0.12, +0.86] | 0.351 |

Hedges' g with bias correction and bootstrap 95% confidence intervals (5,000 resamples), pre-intervention sessions, n = 15, averaged across all 32 channels. Values in the third and fourth columns use the conventional full-band denominator, which is the primary analysis; those in the fifth and sixth exclude delta from total power. Values of q are Benjamini–Hochberg false-discovery-rate corrected within each contrast across the six bands. Movement is the mean of the follow, lead, and improvisation conditions; mirroring is the mean of follow and lead. All values were computed with a common 4-s analysis window. The fast-band effects are essentially unaffected by this choice; the alpha contrasts are the ones that shift, most notably improvisation versus mirroring, which changes sign. No claim in the main text rests on that contrast.

**Table S2. SpecParam model fit quality by condition.**

| Condition | n fits | Mean $r^2$ | Median $r^2$ | % below 0.90 | % below 0.80 | Mean error | Mean exponent |
|---|---|---|---|---|---|---|---|
| Resting baseline | 832 | 0.918 | 0.976 | 24.9 | 10.7 | 0.0461 | 0.916 |
| Eye gaze | 864 | 0.879 | 0.958 | 37.2 | 21.5 | 0.0520 | 0.734 |
| Conversation | 864 | 0.765 | 0.845 | 62.8 | 43.2 | 0.0651 | 0.573 |
| Mirroring (teacher-led) | 864 | 0.807 | 0.891 | 52.2 | 32.2 | 0.0532 | 0.605 |
| Mirroring (participant-led) | 864 | 0.764 | 0.865 | 59.6 | 42.4 | 0.0588 | 0.522 |
| Free improvisation | 864 | 0.778 | 0.870 | 56.5 | 39.7 | 0.0583 | 0.575 |

Values of $r^2$ are the coefficient of determination for the SpecParam model fit. Fixed aperiodic mode, fit over 2–44 Hz, pre- and post-intervention sessions pooled, participants only. Each channel of each recording contributes one fit. Fit quality declines in the active conditions relative to rest, and model fit is positively associated with the estimated exponent across all fits (Spearman rho = +0.78), such that poorer fits return flatter exponents. This association is unchanged by analysis window length (4 s versus condition-specific windows) and by aperiodic mode (fixed versus knee), indicating that it reflects a property of the measure, in that spectra with fewer distinct oscillatory peaks offer the model less structure to fit, rather than a pipeline artifact. Condition effects on the exponent attenuate but persist in direction when restricted to well-fit channels: eye gaze versus baseline gives $g = -0.97$ across all fits, −0.57 at r-squared of at least 0.90, and −0.40 at r-squared of at least 0.95.

**Table S3. Participant flow and analysis samples.**

| Stage | Dance | Control | Withdrew | Total |
|---|---|---|---|---|
| Enrollment and allocation | | | | |
| Consented and assessed | — | — | — | 18 |
| Withdrew following pre-intervention assessment | — | — | 4 | 4 |
| Randomized | 7 | 7 | — | 14 |
| EEG sessions recorded | | | | |
| Pre-intervention | 7 | 6 | 4 | 17 |
| Post-intervention | 7 | 6 | 0 | 13 |
| EEG analyzed (effective rank at least 15) | | | | |
| Pre-intervention | 7 | 5 | 3 | 15 |
| Post-intervention | 7 | 5 | 0 | 12 |
| Paired (both timepoints) | 7 | 4 | 0 | 11 |
| Excluded by effective-rank screening | | | | |
| Pre-intervention (effective rank 2 and 11) | 0 | 1 | 1 | 2 |
| Post-intervention (effective rank 14) | 0 | 1 | 0 | 1 |
| Accelerometry (not subject to rank criterion) | | | | |
| Pre-intervention | 7 | 6 | 4 | 17 |
| Post-intervention | 7 | 6 | 0 | 13 |
| Behavioral outcomes | | | | |
| Completed pre- and post-intervention batteries | 6 | 7 | 0 | 13 |

Counts are participants. Section headings are unindented; the rows beneath each are indented. Withdrew denotes participants who completed the pre-intervention assessment but did not enter the intervention and were not assigned to an arm. Pre-intervention assessment preceded randomization, so participants who subsequently withdrew still contributed a pre-intervention session, and the pre-intervention EEG sample therefore exceeds the randomized sample. Effective-rank screening was applied to the artifact-corrected recordings and governs all spectral, connectivity, and corticokinematic-coherence analyses, which therefore share a single analysis sample. Three participant recordings were excluded: two at pre-intervention (effective rank 2 and 11) and one at post-intervention (effective rank 14). One participant contributed a post-intervention session only and one a pre-intervention session only, which is why the paired sample of 11 is smaller than either timepoint. One post-intervention session lacked the resting-baseline condition and contributed to the five remaining conditions only. Accelerometry is not subject to the rank criterion and retains all recorded sessions. One dance participant completed post-intervention EEG but not the questionnaires; one control participant completed the intervention and questionnaires but never underwent EEG.

**Table S4. Absolute, relative, and aperiodic-adjusted band power across the graded decomposition.**

| Band | Contrast | Absolute g | Relative g | Periodic g |
|---|---|---|---|---|
| Delta | Eye gaze – baseline | +0.30 | -0.31 | +0.99* |
| Delta | Conversation – eye gaze | +1.72* | +0.36 | +0.67* |
| Delta | Movement – eye gaze | +0.97* | -0.07 | +0.17 |
| Delta | Improvisation – mirroring | +0.52 | -0.53 | +0.03 |
| Theta | Eye gaze – baseline | +0.32 | -0.32 | +0.39 |
| Theta | Conversation – eye gaze | +1.63* | -0.01 | -0.64* |
| Theta | Movement – eye gaze | +1.23* | +0.42 | -0.39 |
| Theta | Improvisation – mirroring | +1.00* | +0.78* | +0.01 |
| Alpha | Eye gaze – baseline | -0.13 | -0.87* | -0.90* |
| Alpha | Conversation – eye gaze | +1.08* | -0.47 | -1.24* |
| Alpha | Movement – eye gaze | +0.83* | -0.07 | -0.75* |
| Alpha | Improvisation – mirroring | +0.80* | +0.23 | -0.80* |
| Low beta | Eye gaze – baseline | +0.46 | +0.31 | -0.44 |
| Low beta | Conversation – eye gaze | +1.86* | +0.07 | -1.24* |

| | | | | |
|---|---|---|---|---|
| Low beta | Movement – eye gaze | +1.70* | +0.42 | -0.73* |
| Low beta | Improvisation – mirroring | +1.00* | +0.48 | -0.33 |
| High beta | Eye gaze – baseline | +0.77* | +1.02* | +0.28 |
| High beta | Conversation – eye gaze | +2.02* | +0.51 | -0.45 |
| High beta | Movement – eye gaze | +2.11* | +0.78* | -0.70* |
| High beta | Improvisation – mirroring | +0.85* | +0.38 | -0.43 |
| Gamma | Eye gaze – baseline | +0.81* | +0.95* | +0.57 |
| Gamma | Conversation – eye gaze | +2.12* | +0.91* | +1.08* |
| Gamma | Movement – eye gaze | +2.38* | +1.13* | +0.39 |
| Gamma | Improvisation – mirroring | +0.85* | +0.52 | +0.34 |

Hedges' g with bias correction, pre-intervention sessions, n = 15, averaged across all 32 channels. Absolute power is log-transformed band-averaged power; relative power is each band's share of total power across 1–45 Hz; periodic power is the mean residual between the observed log-power spectrum and the fitted aperiodic component over the 2–44 Hz SpecParam fit range, and is therefore continuous and defined for every channel rather than conditional on detection of a discrete oscillatory peak. Asterisks denote significance after Benjamini–Hochberg false-discovery-rate correction applied within contrast across the six bands, separately for each metric. Movement is the mean of the follow, lead, and improvisation conditions; mirroring is the mean of follow and lead. All values were computed with a common 4-s analysis window. Relative power is not independent of the aperiodic fit: relative gamma and high beta correlate with the aperiodic exponent at rho = −0.95 and −0.94 across condition cells, whereas periodic power is largely independent of it (rho = +0.39, −0.15, and +0.08 for alpha, high beta, and gamma). The alpha reduction at eye gaze is confirmed as an oscillatory change even though absolute alpha is unchanged, because the aperiodic component rises beneath a steady alpha peak; the high-beta increase does not survive aperiodic adjustment, whereas the gamma increase is partly oscillatory.
