## Supplementary Figures for "Social attention and coordinated movement produce dissociable spectral and network changes during dyadic improvisational dance"

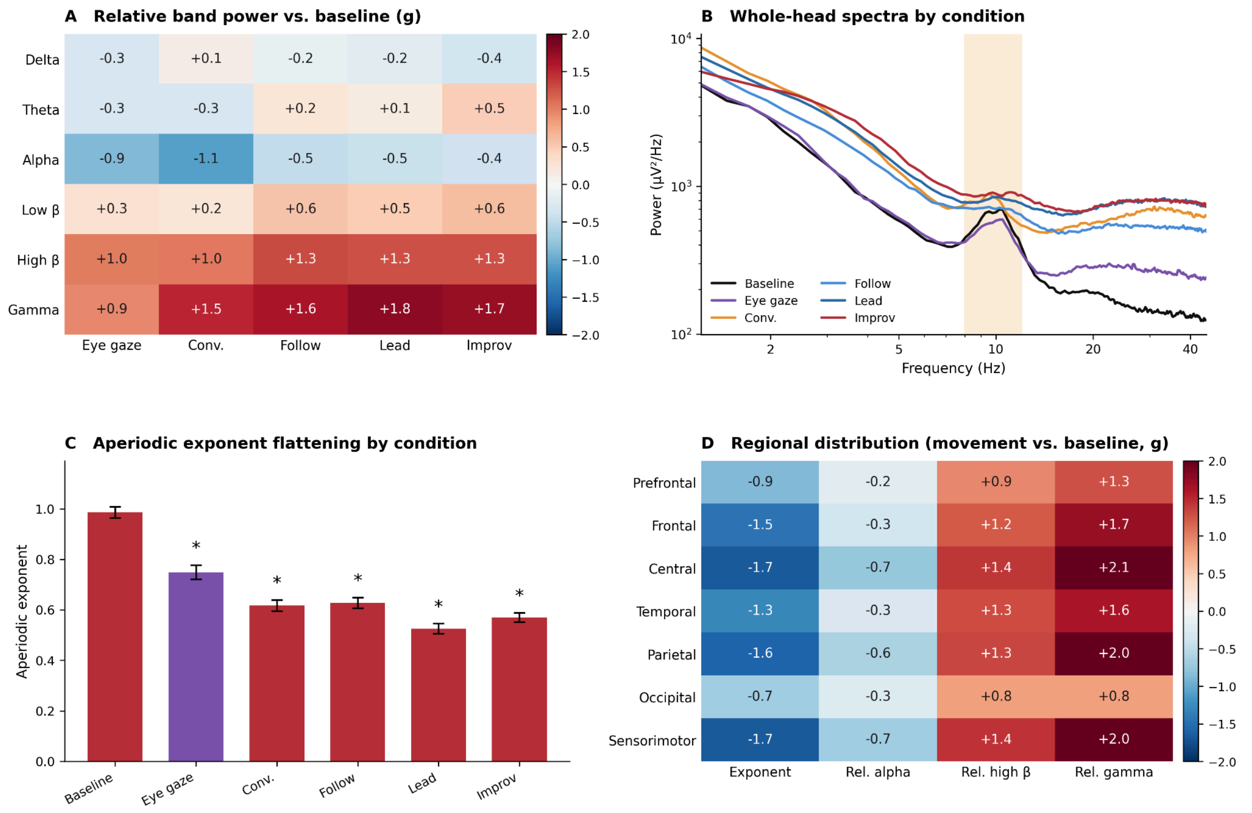


***Supplementary Figure S1. Spectral detail: relative power, aperiodic flattening, and regional distribution.*** *(A) Relative band-power change from baseline (Hedges g) across bands and conditions. (B) Whole-head power spectra by condition, illustrating the aperiodic flattening. (C) Aperiodic exponent by condition, with eye gaze highlighted. (D) Regional distribution of the aperiodic and relative-power effects across seven ROIs, comparing the movement conditions with resting baseline, showing the sensorimotor/centroparietal maximum and occipital minimum. This figure provides the spectral and topographic detail underlying the aperiodic and power measures summarized in Figures 3–4.*

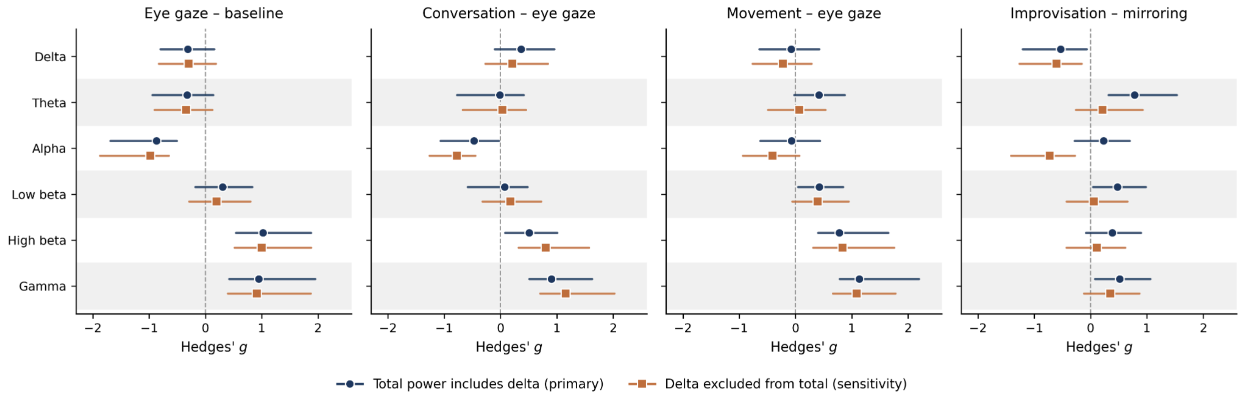


***Supplementary Figure S2. Relative-power effect sizes are robust to the treatment of delta power, except for alpha during movement.*** *Each panel shows one contrast in the graded decomposition. Points are Hedges' g with bootstrap 95% confidence intervals for each of the six frequency bands, computed with the conventional full-band denominator (dark circles, primary analysis) and with delta excluded from total power (orange squares, sensitivity analysis). Fast-band effects (high beta, gamma) are unchanged by the choice of denominator at every contrast. Alpha suppression is larger under the delta-free normalization during the movement contrasts, which is the sole finding materially affected. Pre-intervention sessions, n = 15, all 32 channels.*

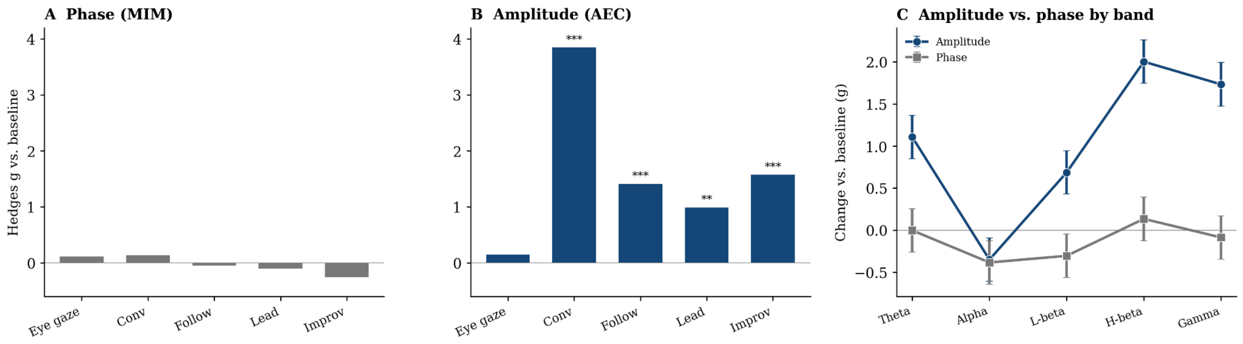


***Supplementary Figure S3. Within-brain connectivity by condition.*** *Whole-brain high-frequency connectivity change from baseline by condition. (A) Phase connectivity (MIM) shows only small condition effects. (B) Amplitude connectivity (AEC) increases, most during conversation and the movement conditions. (C) Frequency-resolved change: amplitude connectivity is concentrated in and rises toward the fast bands while phase connectivity remains flat. This figure provides the by-condition and frequency-resolved connectivity detail underlying Figures 3–4.*

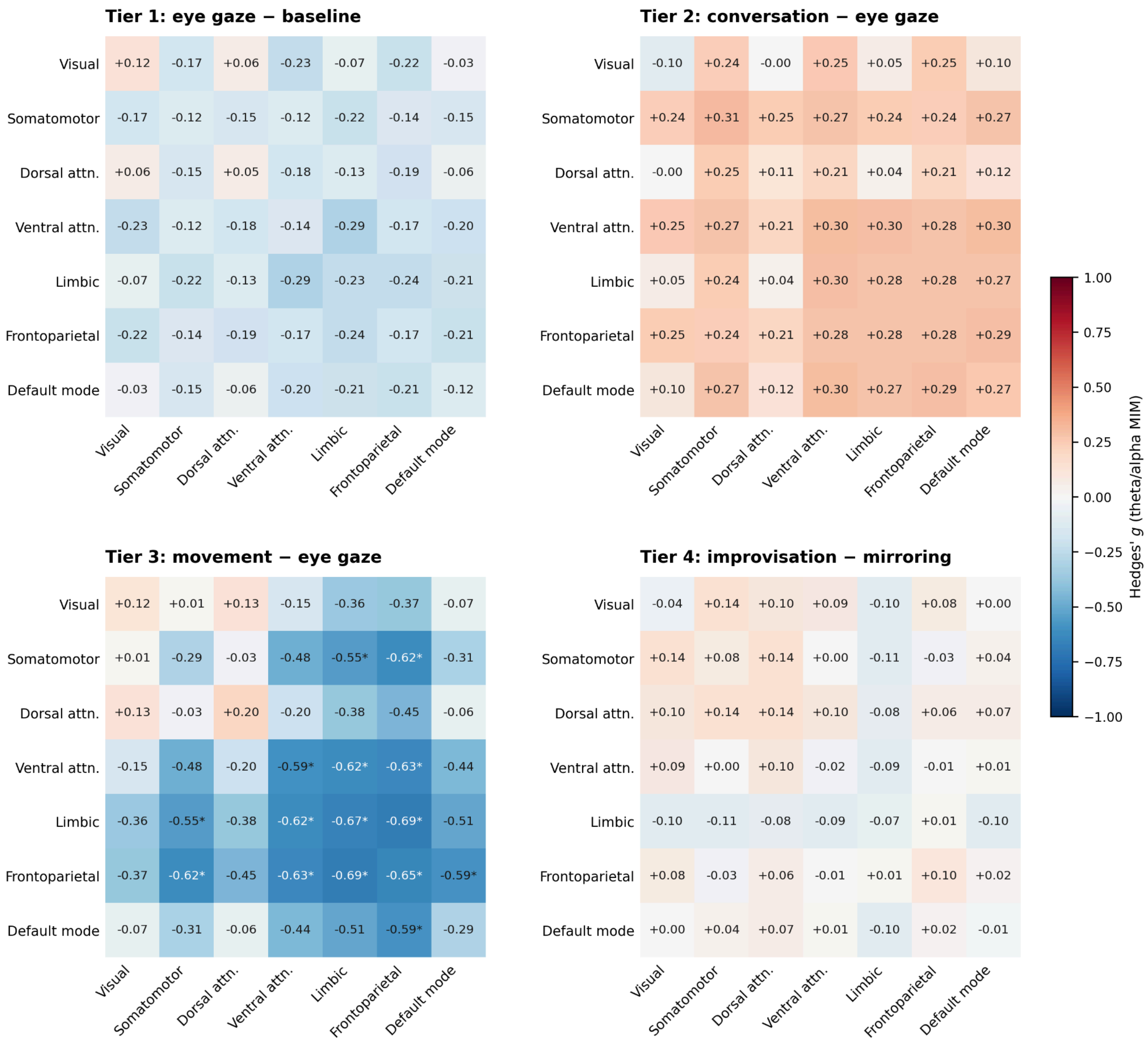


***Supplementary Figure S4. Network matrices for the main condition effects.*** *Network-resolved connectivity change (active conditions vs. baseline) across seven Yeo networks. (A) High-frequency amplitude connectivity (AEC) increases broadly across network pairs. (B) Theta/alpha phase connectivity (MIM) decreases, most in frontoparietal-control connections including DMN–FPN. This figure provides the network topography of the connectivity effects reported for the coordinated-movement tier in Figures 3–4.*
