## Supplementary material for "Social attention and coordinated movement produce dissociable spectral and network changes during dyadic improvisational dance": Manuscript Tables

**Table 1. The graded decomposition of dyadic dance: incremental change in individual-brain measures at each tier.**

| Measure | Eye gaze – baseline | Conversation – eye gaze | Movement – eye gaze | Improvisation – mirroring |
|---|---|---|---|---|
| Aperiodic exponent | -0.97* [-1.94, -0.46] | -0.48 [-1.21, +0.01] | -0.61* [-1.50, -0.11] | -0.04 [-0.56, +0.49] |
| Relative alpha power | -0.87* [-1.69, -0.51] | -0.47 [-1.07, -0.03] | -0.07 [-0.63, +0.43] | +0.23 [-0.29, +0.69] |
| Periodic alpha power | -0.90* [-1.56, -0.49] | -1.24* [-1.77, -0.99] | -0.75* [-1.23, -0.42] | -0.80* [-2.00, -0.24] |
| Relative high-beta power | +1.02* [+0.54, +1.88] | +0.51 [+0.08, +1.01] | +0.78* [+0.40, +1.65] | +0.38 [-0.09, +0.89] |
| Relative gamma power | +0.95* [+0.42, +1.95] | +0.91* [+0.51, +1.63] | +1.13* [+0.78, +2.20] | +0.52 [+0.07, +1.06] |
| Amplitude connectivity (AEC) | +0.15 [-0.37, +0.63] | +2.60* [+1.96, +4.26] | +1.01* [+0.65, +1.86] | +0.64* [+0.18, +1.50] |
| Phase connectivity (MIM) | +0.11 [-0.30, +1.12] | +0.07 [-0.63, +0.49] | -0.50 [-1.21, -0.05] | -0.60 [-1.15, -0.26] |
| DMN–FPN coupling | -0.21 [-0.72, +0.31] | +0.29 [-0.20, +0.70] | -0.59* [-1.09, -0.19] | +0.02 [-0.60, +0.49] |

Values are Hedges' g with bias correction and bootstrap 95% confidence intervals (5,000 resamples), pre-intervention sessions, n = 15. Each tier expresses the incremental contribution of one component of dyadic dance: Tier 1 (eye gaze – baseline) adds nonverbal social attention to solitary rest; Tier 2 adds verbal engagement; Tier 3 adds coordinated whole-body movement; Tier 4 contrasts generative improvisation with structured mirroring. Movement is the mean of the follow, lead, and improvisation conditions; mirroring is the mean of follow and lead. Asterisks denote significance after Benjamini–Hochberg false-discovery-rate correction applied within tier across the eight measures. Spectral measures are channel-level power averaged across 32 channels and the SpecParam aperiodic exponent (2–44 Hz, fixed mode); relative power is each band's share of total power, and periodic alpha power is the mean residual above the fitted aperiodic component, which is independent of the exponent (rho = +0.39) where relative power is not (relative gamma rho = −0.95). Amplitude connectivity (AEC) and phase connectivity (MIM) are source-level whole-brain means across 2,278 region pairs, each averaged over the high-beta and gamma bands. DMN–FPN coupling is the theta/alpha multivariate interaction measure between the default-mode and frontoparietal-control networks. Because the connectivity rows average across two bands, band-resolved effects are not visible here; the gamma-specific amplitude increase and the low-beta and gamma phase decreases at Tier 4, reported in the text, survive correction within their own band families. Tier 3 and Tier 4 contrasts involve conditions that differ in movement vigor.

**Table 2. Socio-emotional and cognitive outcomes before and after the four-week intervention.**

| Measure | Dance pre | Dance post | Control pre | Control post | g | n |
|---|---|---|---|---|---|---|
| MEES Responsive crying | 12.8 (1.5) | 13.2 (1.2) | 12.0 (1.4) | 11.1 (0.7) | +0.82 | 6 / 7 |
| FIAT-Q Interpersonal conflict | 56.6 (15.5) | 50.2 (13.2) | 63.1 (6.0) | 66.1 (7.1) | -1.08 | 5 / 7 |
| PSS Perceived stress | 17.0 (5.2) | 11.8 (5.7) | 19.9 (3.6) | 19.3 (4.5) | -0.99 | 6 / 7 |
| Stroop congruent RT (ms) | 1251 (495) | 1020 (289) | 1221 (678) | 1276 (641) | -0.91 | 6 / 7 |
| SCS Social connectedness | 92.7 (15.3) | 96.0 (15.6) | 78.4 (18.0) | 82.0 (14.8) | -0.02 | 6 / 7 |

Values are mean (SD). Hedges' g compares pre-to-post change between arms, with positive values indicating a greater increase in the dance group; the direction favoring the hypothesis differs by measure, since higher scores are better for MEES and SCS and lower scores are better for FIAT-Q, PSS, and Stroop reaction time. Sample sizes differ by measure because of item-level missingness and are given as dance / control. After Benjamini–Hochberg correction within scale, one effect survived: MEES responsive crying (ANCOVA controlling for baseline, $p = 0.006$, partial eta-squared = 0.54, corrected $p = 0.044$), on which the dance group increased while the control group decreased. Effects for interpersonal conflict, perceived stress, and Stroop congruent reaction time were large and in the hypothesized direction but did not survive correction. All intervention contrasts are underpowered at this sample size and are reported as pilot effect-size estimates rather than as tests of efficacy. Stroop reaction time was computed from correct trials at pre-intervention and from all trials at post-intervention.
